# Somatic mtDNA Mutation Burden Shapes Metabolic Plasticity in Leukemogenesis

**DOI:** 10.1101/2024.09.25.615001

**Authors:** Xiujie Li-Harms, Jingjun Lu, Yu Fukuda, John Lynch, Aditya Sheth, Gautam Pareek, Marcin Kaminski, Hailey Ross, Christopher W. Wright, Huiyun Wu, Yong-Dong Wang, Geoff Neal, Amber Smith, Peter Vogel, Stanley Pounds, John Schuetz, Min Ni, Mondira Kundu

## Abstract

The role of somatic mitochondrial DNA (mtDNA) mutations in leukemogenesis remains poorly characterized. To determine the impact of somatic mtDNA mutations on the process, we assessed the leukemogenic potential of hematopoietic progenitor cells (HPCs) from mtDNA mutator mice (Polg D257A) with or without NMyc overexpression. We observed a higher incidence of spontaneous leukemogenesis in recipients transplanted with heterozygous Polg HPCs and a lower incidence of NMyc-driven leukemia in those with homozygous Polg HPCs compared to controls. Although mtDNA mutations in heterozygous and homozygous HPCs caused similar baseline impairments in mitochondrial function, only heterozygous HPCs responded to and supported altered metabolic demands associated with NMyc overexpression. Homozygous HPCs showed altered glucose utilization with pyruvate dehydrogenase inhibition due to increased phosphorylation, exacerbated by NMyc overexpression. The impaired growth of NMyc-expressing homozygous HPCs was partially rescued by inhibiting pyruvate dehydrogenase kinase, highlighting a relationship between mtDNA mutation burden and metabolic plasticity in leukemogenesis.

**TEASER:** Somatic mtDNA mutations as drivers of metabolic change in the development of leukemia.

## INTRODUCTION

Metabolic plasticity, defined as the ability of cells to adapt their metabolic pathways in response to environmental and genetic changes, is critical for the development and progression of tumors (*1, 2*). This adaptability allows cancer cells to meet their increased energy demands and survive under adverse conditions. Historically, research elucidating the mechanisms by which genomic changes contribute to tumorigenesis has primarily focused on mutations in genes encoded by nuclear DNA (nDNA). However, given the central role of mitochondrial metabolism in tumorigenesis and the potential for mutations in the mitochondrial genome to influence metabolic plasticity, there is a growing interest in understanding the contribution of mitochondrial DNA (mtDNA) mutations during tumorigenesis and determining whether targeting mitochondrial metabolism holds therapeutic potential (*3–6*)

The 16-kb circular mitochondrial genome encodes 37 genes, including 13 polypeptides as well as the 2 rRNAs and 22 tRNAs required for their translation (*7*). These mtDNA-encoded proteins assemble with more than 80 nDNA-encoded proteins to form the mitochondrial respiratory chain complexes that drive oxidative phosphorylation (OXPHOS) (*5, 8, 9*). The importance of the mtDNA-encoded OXPHOS components in oncogenesis is highlighted by the observation that cells devoid of mtDNA (i.e., rho-0 cells) lack tumorigenic potential, which can be restored in such cells through the transfer or acquisition of normal mitochondria (*10, 11*).

As with nDNA, mutations in mtDNA arise due to defects in nucleotide metabolism and replicative infidelity (*12*). The first report of an mtDNA mutation in a renal oncocytoma was published decades ago (*13*), followed by the discovery of somatic mtDNA mutations in human malignancies such as colorectal cancer (*14*) and acute leukemia(*15, 16*). More recent large-scale analyses of mtDNA variants in human cancers have revealed that approximately 60% of adult cancers harbor somatic mtDNA variants with a wide range of allele frequencies (*17, 18*). Although pathogenic mtDNA mutations clearly influence cellular energy production and redox status when present at high levels (i.e., >50%), these mtDNA mutations can also engage mitochondrial–nuclear cross-talk pathways and elicit epigenetic and transcriptional changes when present at allele frequencies as low as 20%–30% (*19*), levels similar to those typically observed in primary tumor samples (*17*). Although some studies have suggested that the ability of mtDNA mutations to alter the growth of established tumors depends on the pathogenicity of the mutation and level of heteroplasmy (*7, 20*), the effect of mtDNA mutations and their cumulative burden on tumor initiation and progression remains unclear due to the shortage of model systems.

MtDNA mutator mice express an exonuclease-inactive polymerase mutant (*Polg^D257A^*, referred to hereafter as *Polg^mut^*) with impaired proofreading capacity. The consequent accumulation of somatic mtDNA mutations and associated mitochondrial respiratory chain dysfunction contribute to the shortened lifespans due to accelerated onset of ageing-related disorders (e.g., myelodysplastic syndrome-like disease, osteoporosis) in homozygous (Hom; *Polg^mut/mut^*) mtDNA mutator mice (*21–23*). Heterozygous (Het; *Polg^wt/mut^*) mtDNA mutator mice have normal lifespans, but show an increased incidence of indolent B-cell lymphomas (*24*). The increased frequency of somatic mtDNA mutations across wildtype (WT; *Polg^wt/wt^*), Het, and Hom mtDNA mutator mice provides a unique system to evaluate the consequences of varying the mtDNA mutation burden on tumor development.

In the current study, we sought to investigate the effect of varying the mtDNA mutation burden on the development of acute leukemia induced in mice by overexpressing NMyc in hematopoietic progenitor cells (HPCs) (*25*). NMyc is a member of the MYC family of transcription factors, whose deregulated expression in a variety of cancers, including hematopoietic malignancies, is associated with extensive rewiring of metabolic and energetic pathways (*26–29*). Our results suggest that the degree of metabolic plasticity, influenced by the burden of mtDNA mutations, plays a critical role in determining the tumorigenic potential of HPCs.

## RESULTS

### Higher incidence of spontaneous leukemia but comparable incidence of NMyc-driven leukemia derived from Het HPCs compared to WT HPCs

Whole mitochondrial genome sequencing (*30*) was performed on DNA extracted from bone marrow (BM)-derived cells isolated from WT (*Polg*^wt./wt.^), Het (*Polg^wt/mut^*), and Hom (*Polg^mut^*^/*mut*^) mice to determine the relationship between the frequency of somatic mtDNA mutations and mutant Polg allele dosage. This confirmed the expected increase in the number of somatic mtDNA mutations at very low variant allele fractions (i.e., 1%) in Het and Hom BM cells compared to WT (**Fig. S1A)** (*31, 32*).

We then isolated HPCs from 2- to 5-month-old WT, Het, and Hom mice, and transduced them with either a retroviral construct expressing YFP alone (MSCV-IRES-YFP) as a control (Ctrl) or a retroviral construct expressing NMyc and YFP (MSCV-NMyc-HA-IRES-YFP). The transduced HPCs were transplanted into lethally irradiated syngeneic recipients to evaluate their leukemogenic potential *in vivo* (**Fig. 1A**). Flow cytometric analyses showed comparable percentages of transduced YFP+ cells among the recipient groups prior to and shortly after transplantation (**Fig. S1B** and **C**).

**Figure 1.**
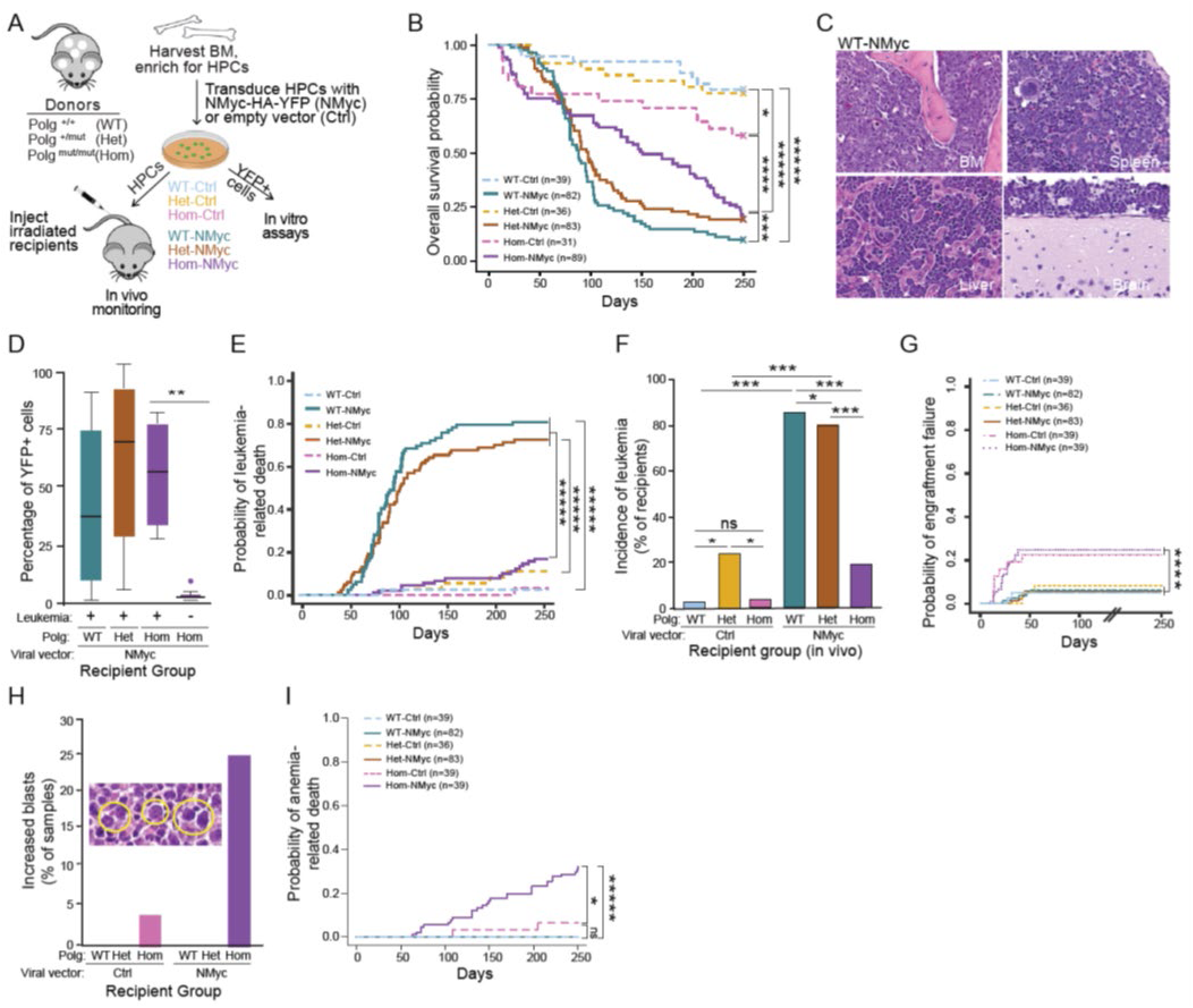
Effect of mtDNA mutation burden on leukemogenic potential. **(A)** Schematic representation of experimental setup. Hematopoietic progenitor cells (HPCs) were isolated from bone marrow (BM) of 2- to 5-month-old homozygous (Hom, *Polg^mut/mut^*) and heterozygous (Het, *Polg^wt/mut^*) mtDNA mutator mice, as well as wildtype (WT, *Polg*^wt/wt^) littermate controls. These HPCs were transduced with either MSCV-NMyc-HA-IRES-YFP (NMyc) or control MSCV-IRES-YFP (Ctrl) retrovirus. Transduced HPCs were subsequently transplanted into lethally irradiated C57BL/6 syngeneic recipients for *in vivo* assessment of leukemogenic potential *o*r sorted and evaluated *in vitro* analysis. **(B)** Kaplan-Meier survival curves for recipients transplanted with different HPC populations: WT-Ctrl (n=39), WT-NMyc (n=82), Het-Ctrl (n=36), Het-NMyc (n=83), Hom-Ctrl (n=31), and Hom-NMyc (n=89). A log rank test was conducted to examine if the survival distributions differed among the recipient groups. Given that there was a significant overall test result (p<0.0001) suggesting the survival distributions were different among the 6 genetic groups, we further examined which 2 groups were significantly different by performing a pairwise comparison between the genetic groups. The p values of individual comparisons are shown in the figure: *P<0.05; ***P<0.005; ****P<0.001; *****P<0.0001. **(C)** Representative hematoxylin and eosin (HE) stained sections show typical leukemic infiltrates in the BM, spleen, liver, and brain of a WT-NMyc recipient. **(D)** Levels of circulating YFP+ leukocytes were analyzed by flow cytometry using peripheral blood collected from recipients transplanted with NMyc-expressing HPCs as they became symptomatic. Hom-NMyc mice were later separated into two groups (leukemic and non-leukemic) based on necropsy results. Sample sizes: WT-NMyc (n=27), Het-NMyc (n=21), Hom-NMyc (tumor cases, n=4), and Hom-NMyc (non-tumor cases, n=8); **P<0.01 (Student’s *t*-test). **(E)** The Fine and Gray model was used to analyze the probability of death by various causes using subdistribution analysis of competing risks. Plots show cumulative incidence functions (CIF) for leukemia-related deaths. Gray’s test for equality was used to compare CIFs: ****P<0.001. **(F)** Graph showing incidence of leukemia in recipients: WT-Ctrl (n=1/32), WT-NMyc (n=68/77), Het-Ctrl (n=6/25), Het-NMyc (n=58/75), Hom-Ctrl (n=1/26), Hom-NMyc (n=16/84). *P<0.05; ***P<0.005; ns: no significant difference. **(G)** Plots showing CIF for deaths related to engraftment failure. Gray’s test for equality was used to compare CIFs: ****P<0.001. **(H)** Increased numbers of scattered blasts (highlighted by yellow circles) were identified in H&E-stained BM and spleen samples from mice transplanted with Hom HPCs (NMyc>Ctrl). Graph shows the percentage of cases bearing increased numbers of blasts from each group: WT-Ctrl (n=0/32), WT-NMyc (n=0/77), Het-Ctrl (n=0/25), Het-NMyc (n=0/75), Hom-Ctrl (n=2/26), Hom-NMyc (n=37/84). **(I)** Plots show CIFs for anemia-related deaths. Gray’s test for equality was used to compare CIFs: *P<0.05; *****P<0.0001.

Transplant recipients were closely monitored and sacrificed upon reaching humane endpoints to calculate overall survival probability (**Fig. 1B**). Necropsies were performed to determine the cause of death and for histologic and immunophenotypic evaluation of tissue infiltrates (**Table S1**). Transplantation of NMyc-expressing WT HPCs resulted in a highly penetrant phenotype in recipients (hereafter WT-NMyc mice), characterized by infiltration of the spleen, liver, brain, and BM with leukemic blasts (**Fig. 1C**), consistent with previous observations (*25*). Complete blood counts taken from mice that reached a humane endpoint revealed an increase in white blood cells (WBCs) (**Fig. S1D**) and reduced platelets (**Fig. S1E**), but no significant differences in red blood cell (RBC) counts (**Fig. S1F**) or hemoglobin (Hgb) concentration (**Fig. S1G**) in WT-NMyc mice compared to WT-Ctrl mice. Flow cytometric analyses of peripheral blood revealed high levels of circulating YFP+ cells in end-stage WT-NMyc mice (**Fig. 1D**), supporting the diagnosis of leukemia. Immunohistochemical analyses revealed extensive infiltration of the BM and other organs by hemagglutinin (HA)+ leukemic blasts expressing T-cell, B-cell, and/or myeloid markers (**Fig. S1H**). A combination of flow cytometry and immunohistochemical analyses was used to classify the disease in each recipient as T-cell, B-cell, myeloid or mixed lineage leukemia (**Fig. S1I**). The probability of leukemia-associated mortality peaked around 160 days post-transplantation among WT-NMyc mice (**Fig. 1E**), with over 80% developing acute leukemia (**Fig. 1F**).

Mice transplanted with Het-NMyc HPCs (hereafter Het-NMyc mice) developed leukemia and reached humane endpoints at a rate similar to their WT counterparts (**Fig. 1B, D**, and **F** and **S1D–I**). Notably, a significantly higher proportion of mice transplanted with Het-Ctrl HPCs developed acute leukemia by the end of the study than those transplanted with WT-Ctrl HPCs (∼20% vs <3%, P=0.032) (**Fig. 1F**). These results suggest that Het HPCs could support NMyc-induced leukemia at a rate similar to WT HPCs but were more susceptible to spontaneous leukemia development than WT HPCs.

### Lower incidence of NMyc-induced leukemia derived from Hom HPCs than WT or Het HPCs

We next assessed the fate of Hom HPCs in transplant recipients. There was a small but significant (P<0.001) increase in the incidence of early mortality (by 2 months post-transplantation) in recipients transplanted with either control or NMyc-expressing Hom HPCs (hereafter Hom-Ctrl or Hom-NMyc mice) (**Fig. 1B**) due to engraftment failure (**Fig. 1G**). There were no significant differences in peripheral blood counts between successfully engrafted Hom-Ctrl mice and WT or Het-Ctrl mice (**Fig. S1D–G**). Although the overall survival probability of Hom-NMyc mice was significantly lower than those in the Hom-Ctrl group (P<0.001) (**Fig. 1B**), less than 20% of the Hom-NMyc mice showed hematologic or histopathologic features consistent with acute leukemia at the end stage (**Fig. 1E and F**). Accordingly, the incidence of leukemia and the leukemia-associated mortality of Hom-NMyc mice were significantly lower than those in WT-NMyc or Het-NMyc groups (P<0.005) (**Fig. 1E and F**). An additional 25% of the Hom-NMyc mice showed increased numbers of scattered blasts in their BM (**Fig. 1H**). Regardless of the presence of acute leukemia, RBC counts and Hgb concentrations in Hom-NMyc mice were significantly lower than those in all other groups of mice (P<0.001) including the Hom-Ctrl mice (**Fig. S1F and G**). The increased incidence of anemia and probability of anemia-related death in these mice (**Fig. 1I**) were similar to those observed in aged Hom mtDNA mutator mice in previous studies (*21, 22, 33–35*). Therefore, although the upregulation of NMyc in Hom HPCs appeared to accelerate the dysregulation of erythroid differentiation, leading to the onset of anemia in recipients, the increased burden of somatic mtDNA mutations in Hom HPCs impaired the leukemogenic potential of NMyc.

### Alterations in mtDNA mutation burden, mtDNA copy number, and common oncogenic mutations in leukemic cells from Hom mice

We next compared the mtDNA mutation burden and copy number in bulk BM cells derived from WT, Het, and Hom mice, with those from leukemia-infiltrated recipients (i.e., WT-NMyc, Het-NMyc, and Hom-NMyc mice). Sequencing entire mitochondrial genomes showed significant increases in the overall burden of somatic mtDNA mutations in the Het and Hom leukemia samples compared to WT leukemia samples and all control BM samples, regardless of genotype (P<0.05) (**Fig. 2A and B**). Using mtDNA sequencing data, we also examined the ratio of non-synonymous to synonymous (dN/dS) substitutions in protein-coding genes among normal and leukemia-infiltrated BM samples. The dN/dS ratio is a measure used to compare the rates of mutations that alter the amino acid sequence of a protein (non-synonymous) to those that do not (synonymous) and is useful for understanding selective pressures on genes. A higher dN/dS ratio was observed in the leukemic samples compared to the control BM samples (**Fig. 2C**), particularly in the Het-NMyc and Hom-NMyc leukemic samples at low-to-intermediate variant allele fractions (0.2 to 0.5). These findings suggest that intermediate levels of non-synonymous somatic mtDNA mutations were enriched, or at least more tolerated, in leukemic cells than in normal BM cells.

**Figure 2.**
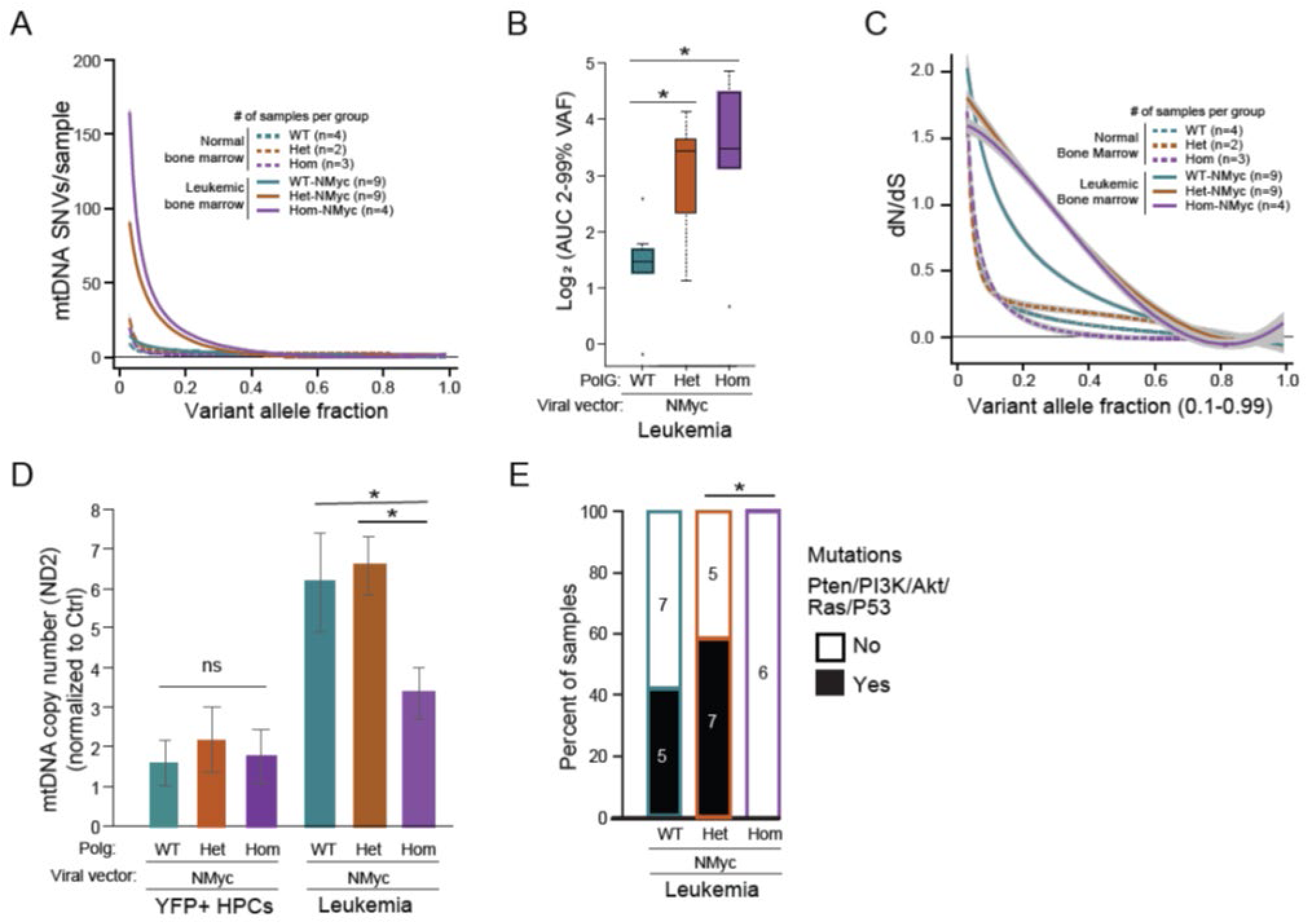
Analysis of mtDNA mutation burden, copy number, and common oncogenic mutations in leukemic samples. **(A)** Comparison of mtDNA mutation burden in cells derived from normal and leukemia-infiltrated BM samples. Sample size: normal BM - WT (n=4), Het (n=2), and Hom (n=3); leukemic BM - WT-NMyc (n=9), Het-NMyc (n=9), and Hom-NMyc (n=4). **(B)** Analysis of variant allele fractions (VAF) in cells derived from leukemia-infiltrated BM samples. Plots show area under the curve (AUC) values from VAFs ranging from 0.2 to 0.99. Sample sizes: WT-NMyc (n=9), Het-NMyc (n=9), and Hom-NMyc (n=4); *P <0.05. **(C)** The ratio of non-synonymous (dN) to synonymous (dS) in normal and leukemic BM. Sample sizes: normal BM - WT (n=4), Het (n=2), and Hom (n=3); leukemic BM - WT-NMyc (n=9), Het-NMyc (n=9), and Hom-NMyc (n=4). **(D)** Quantification of mtDNA content in YFP+ HPCs prior to transplantation and in cells from leukemia-infiltrated BM samples. The graph shows relative mtDNA copy number (mean ± s.e.m) calculated using ND2 as the mtDNA marker and 18S and the nDNA marker. Ratios were normalized to a control sample that was included in all experiments to allow comparison of values across experiments. Sample size: YFP+ HPCs prior to transplantation - WT-NMyc (n=7), Het-NMyc (n=6), and Hom-NMyc (n=7); leukemic BM - WT-NMyc (n=15), Het-NMyc (n=17), and Hom-NMyc (n=8). *P<0.05 (ANOVA). **(E)** The exome sequences in leukemia-infiltrated BM samples were examined to detect secondary mutations (Pten/PI3K/Akt/Ras/P53) in different groups: WT-NMyc (n=12), Het-NMyc (n=12), and Hom-NMyc (n=6). *P<0.05 (Fisher’s Exact Test).

Assessment of mtDNA copy number via a quantitative PCR approach using primers targeting 100-bp fragments in MT-ND2 and MT-CO2 revealed a significant (P<0.05) decrease in mtDNA content in leukemic cells from Hom-NMyc mice compared to those from WT- or Het-NMyc mice (**Fig. 2D**). No significant differences in the MT-ND2 to MT-CO2 ratio were detected among the samples (all p values >0.3439, Student’s *t-*test) (**Fig. S2**). As MT-ND2 lies within a region commonly deleted in mtDNA mutator mice while MT-CO2 lies outside this region (*22*), these results indicate that the reduced copy number in leukemia samples from Hom-NMyc mice was not an artifact of commonly deleted regions. Notably, no significant differences in mtDNA content were observed among NMyc-expressing HPCs regardless of genotype prior to transplantation (**Fig. 2D**), suggesting that the differences in mtDNA copy number occurred during the leukemic transformation process.

We also performed exome sequencing on leukemia-infiltrated BM samples from WT-NMyc, Het-NMyc, and Hom-NMyc mice to determine if there were any differences in the profile of secondary mutations. Although no mutations were uniquely enriched in the Hom-NMyc samples, mutations affecting the PTEN/PI3K/AKT, RAS, and p53 pathways, which are common in acute leukemias (*36–39*), were identified in 40%–60% of WT-NMyc and Het-NMyc samples (**Fig. 2E**). Together, these findings highlight an association between the impaired transformation potential of Hom-NMyc HPCs and their failure to increase mtDNA content or to acquire additional oncogenic mutations. These defects are not observed in Het-NMyc HPCs despite the intermediate burden of mtDNA mutations, suggesting that HPCs can tolerate a certain burden of mtDNA mutations before they impair leukemogenesis.

### *In vitro* recapitulation of leukemogenic potential via serial passaging in methylcellulose

To determine whether the effect of somatic mtDNA mutations on leukemic transformation could be recapitulated *in vitro*, we examined the growth of sorted YFP+ HPCs in methylcellulose cultures for up to six passages. Consistent with previous studies showing that NMyc-expressing HPCs form colonies of transformed progenitors capable of inducing highly penetrant leukemia upon serial passaging (>3 passages) (*25*), cell and colony counts remained high in WT-NMyc samples through MCP#6, but decreased significantly in WT-Ctrl samples by MCP#4 (**Fig. 3A** and **Fig. S3A–C**). Both the colony and cell count in Hom-NMyc samples were reduced relative to WT-NMyc samples at MCP#4 and MCP#6 (**Fig. 3A and B** and **Fig. S3A–C**), with only a few (2 of 9) samples showing colonies at MCP#6 **(Fig. 3C**). Although the mean colony and cell counts were also reduced in Het-NMyc samples (**Fig. 3A and B** and **Fig. S3A–C**), the percentage of samples with at least one colony at MCP#6 was comparable between WT and Het samples (**Fig. 3C**). Curiously, one Het-YFP sample had colonies at MCP#6 (**Fig. 3C**). Thus, when considering the persistence of colonies at MCP >3 as an indication of leukemogenic potential, the impact of the somatic mtDNA mutations showed similar trends *in vivo* (**Fig. 1F**) and *in vitro* (**Fig. 3C**), with an intermediate burden of mtDNA mutations slightly increasing the potential for spontaneous transformation and a high burden impairing NMyc-induced transformation.

**Figure 3.**
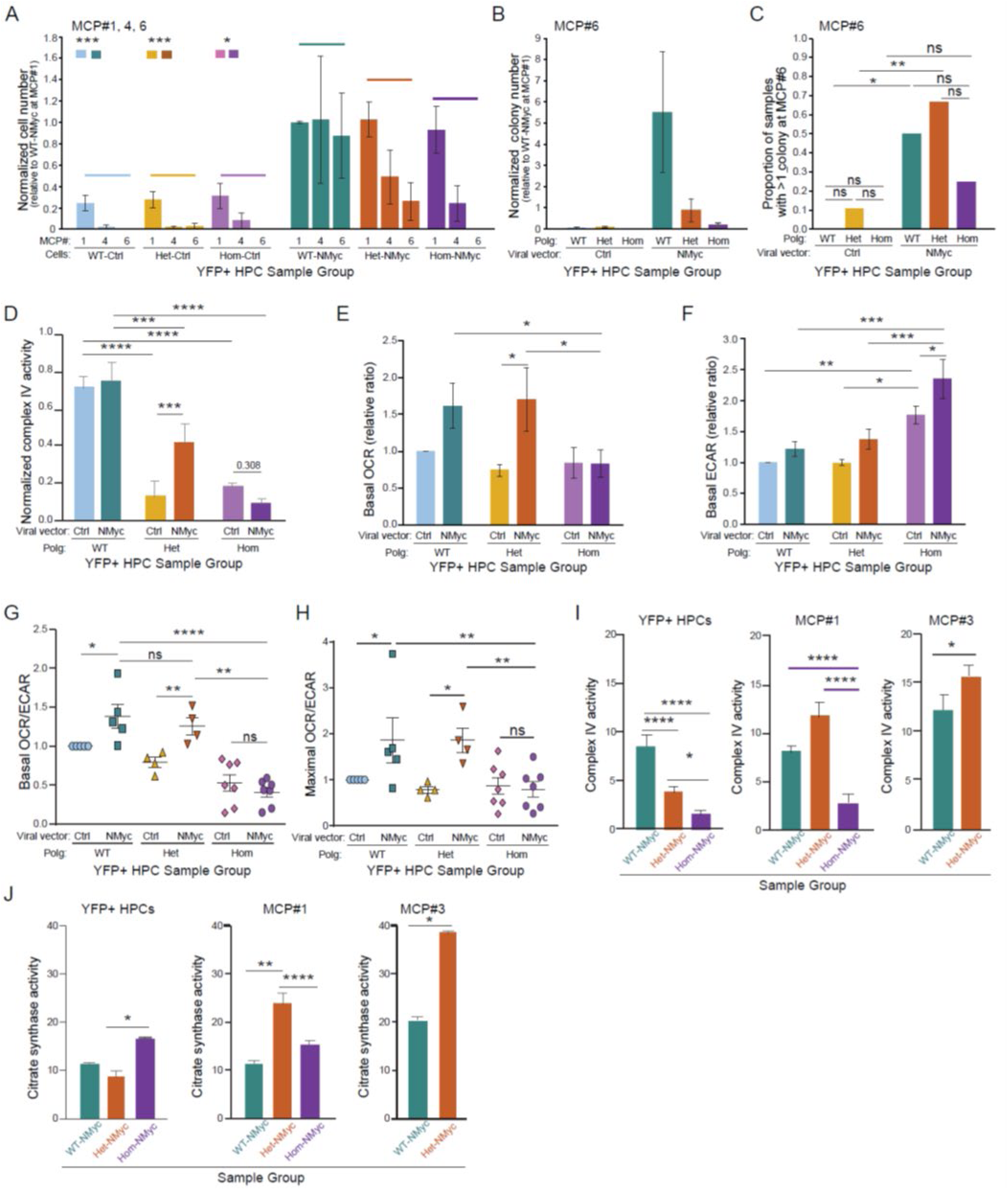
Unlike NMyc-expressing HPCs from Het mtDNA mutator mice, Hom mice fail to compensate for the loss of respiratory complex activity. **(A)** Cell numbers were assessed in serial methylcellulose culture passages (MCP#) at MCP#1, MCP#4, and MCP#6 across different groups and then normalized to the number of WT-NMyc cells at MCP#1. Bar graph shows normalized cell numbers (mean ± s.e.m.). n=8-14 biologic replicates per group. *P<0.05; ***P<0.005 (Student’s *t-*test). **(B)** Colony numbers were assessed at MCP#6 and then normalized to the number of WT-NMyc colonies at MCP#1. Bar graph shows normalized colony numbers (mean ± s.e.m.). n=8-9 biologic replicates per group. **(C)** Bar graph shows the proportion of methylcellulose cultures containing at least one colony at MCP#6 for each group: WT-Ctrl (n=0/8), WT-NMyc (n=4/8), Het-Ctrl (n=1/9), Het-NMyc (n=6/9), Hom-YFP (n=0/8), and Hom-NMyc (n=2/8) *P<0.05; **P<0.01 (Student’s *t-*test). **(D)** Bar graph shows the ratio of complex IV to citrate synthase activity (“Normalized complex IV activity”) using sorted HPCs (pooled from two mice for each genotype). Results from a representative experiment (n=4 technical replicates for each group) are shown. ***P<0.005, ****P<0.001. (one factor ANOVA). **(E-H)** Seahorse assays were used to determine oxygen consumption rates (OCR) and extracellular acidification rates (ECAR) in sorted HPCs. Bar graphs show basal OCR **(E)**, basal ECAR **(F)**, the ratio of basal OCR to ECAR **(G)**, and maximal OCR to ECAR **(H)**. Values were normalized to WT-Ctrl. n=4-7 biological replicates per group. *P<0.05; **P<0.01; ***P<0.005 (one factor ANOVA). **(I, J)** Complex IV **(I)** and citrate synthase **(J)** activities were measured in WT-, Het-, and Hom-NMyc HPCs at MCP#0, MCP#1, and MCP#3. *P<0.05; **P<0.01; ****P<0.001 (one factor ANOVA).

### HPCs from Hom (but not Het) mtDNA mutator mice fail to support the NMyc-induced increase in oxidative metabolism

NMyc expression in leukemic precursors is associated with an increase in oxidative respiration, which is required for oncogenic transformation (*40*). Given the fidelity of the *in vitro* model in recapitulating the pleiotropic effects of somatic mtDNA mutations on NMyc-induced leukemogenesis, we utilized this system to explore the relationships between mtDNA mutations, metabolic responses to NMyc and leukemogenic potential.

Enzymatic activity assays performed on extracts prepared from HPCs (prior to passaging in methylcellulose) revealed that NMyc expression increased overall activity levels of OXPHOS complex IV (CIV) in both WT and Het HPCs, but not in Hom HPCs (**Fig. S3D**). Conversely, citrate synthase (CS) activity increased in response to NMyc expression in all HPCs, regardless of genotype (**Fig. S3E**). Although the ratio of CIV to CS activity, an indicator of mitochondrial function, was not significantly altered by NMyc expression in WT and Hom HPCs, it was significantly increased in Het HPCs (P<0.005; **Fig. 3D**). The NMyc-associated increase in overall CIV activity and the CIV/CS ratio in Het, but not in Hom, HPCs, was notable given that both measures were similarly and significantly (P<0.005 and P<0.001, respectively) lower in Het- and Hom-Ctrl HPCs than those in WT-Ctrl HPCs (**Fig. 3D** and **Fig. S3D**). These findings provided the first indication of increased metabolic plasticity in Het HPCs and reduced metabolic plasticity in Hom HPCs.

Additionally, we measured oxygen consumption rates (OCRs) and extracellular acidification rates (ECARs), indicators of mitochondrial respiration and glycolysis, which are often altered in cells with mitochondrial dysfunction (*41, 42*). Basal OCRs were comparable across all Ctrl samples regardless of genotype, which, given the decreased CIV activity levels in Het- and Hom-Ctrl HPCs (**Fig. 3E**), suggests a reserve of active CIV in HPCs. Nevertheless, the NMyc-induced increase in OCRs in WT and Het HPCs were not detected in Hom HPCs, leading to the lower OCRs in Hom-NMyc HPCs compared to WT-NMyc or Het-NMyc HPCs (P < 0.05, **Fig. 3E**).

The ECARs of Hom HPCs were significantly higher than those in WT and Het HPCs across both the Ctrl and NMyc groups (P<0.005; **Fig. 3F**). This increase was also visibly reflected by the noticeable changes in media color from Hom HPC cultures, associated with a significant elevation in extracellular lactate levels in both Ctrl and NMyc-expressing Hom HPCs (**Fig. S3F, G**). By calculating the OCR to ECAR ratio, an indicator of the cellular preference for oxidative phosphorylation (OXPHOS) over glycolysis (*43*), we found that NMyc expression caused increases in both the basal and maximal OCR:ECAR ratios in WT and Het HPCs, but not in Hom HPCs (**Fig. 3G and H**). This suggests that the higher burden of somatic mtDNA mutations in Hom HPCs impaired OXPHOS activity, making the cells more reliant on aerobic glycolysis. In contrast, Het HPCs retain their mitochondrial oxidative capacity, which is required for the NMyc-induced metabolic adaptation in HPCs.

To determine whether the differences in metabolic response to NMyc expression might contribute to the varying transformation potential in HPCs, we evaluated the activity and protein levels of OXPHOS complexes in NMyc-expressing cells across multiple passages in serial replating experiments. First, we observed that both WT- and Het-NMyc HPCs maintained their CIV and CS activities during passaging, with these activities being more significantly enhanced in NMyc-expressing Het than Hom HPCs (**Fig. 3I and J**). Second, throughout serial passaging, Het Nmyc HPCs showed a significant increase in both CIV and CS activities compared to WT Nmyc HPCs (**Fig. 3I and J**). Nevertheless, in keeping with the mtDNA mutation associated mitochondrial dysfunction in the Het HPCs, the CIV:CS ratio was consistently lower in Het Nmyc HPCs across passages and in the derived leukemic cells (**Fig. S3H, I**). Significant increases in the levels of the mitochondrial marker Tom20 (P<0.005), and OXPHOS complexes II (P<0.05) and V (P<0.05) were observed in leukemic samples derived from Het HPCs compared to WT HPCs (**Fig. S3J-K**), suggesting an increase in mitochondrial content. In contrast, levels of complex I (CI), the OXPHOS complex with the highest proportion of mtDNA-encoded subunits, were significantly lower in leukemic samples derived from Het-NMyc HPCs than in those derived from WT-NMyc HPCs (P=0.01) (**Fig. S3J and K**).

Our findings suggest that functional OXPHOS in the mitochondria of HPCs is required for NMyc-driven leukemic transformation. Moreover, NMyc upregulates mitochondrial oxidative metabolism to support the survival and growth of both HPCs and leukemic cells. Consequently, the higher burden of mtDNA mutations in HPCs from Hom mtDNA mutator mice that limits mitochondrial oxidative capacity and metabolism, reduces the metabolic plasticity needed for HPCs to adjust to increased energy or biosynthetic demands during leukemic transformation and progression. By contrast, although the intermediate burden of mtDNA mutations in the Het HPCs reduces the efficiency of OXPHOS, these HPCs are able to compensate and adapt to the NMyc-induced upregulation of OXPHOS-related mitochondrial functions that occurs during the leukemic transformation process.

### NMyc expression and accumulation of somatic mtDNA mutations are associated with changes in gene expression

To better understand how the accumulation of somatic mtDNA mutations in HPCs influences cellular responses to NMyc expression, we profiled the expression of nDNA-encoded mRNAs isolated from Ctrl and NMyc-expressing WT, Het, and Hom HPCs. We performed a pathway analysis of variance (ANOVA) using Hallmark, and Gene Ontology (GO) gene sets from the Molecular Signatures Database (**Table S2)** (*44–46*). Hierarchical clustering of the Hallmark pathways with differential expression (false discovery rate <0.05) highlighted five distinct response patterns, with three clusters (Clusters 1–3) and two outliers (glycolysis and mitotic spindle pathways) (**Fig. 4A**).

**Figure 4.**
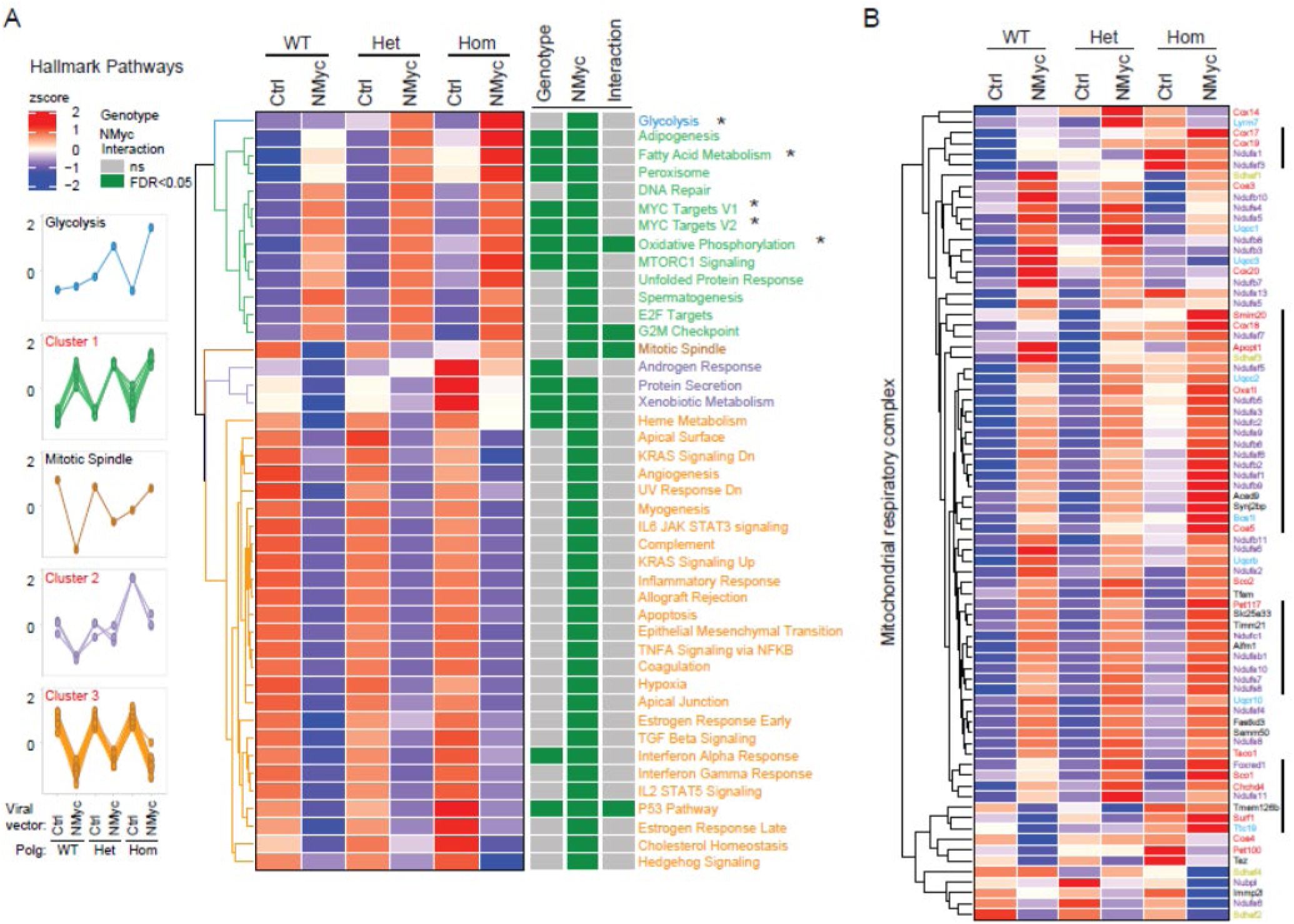
NMyc expression and accumulation of somatic mtDNA mutations are associated with changes in gene expression. **(A)** Heatmap and hierarchical clustering of Hallmark pathways with differential expression shows five distinct patterns of response among the 6 groups of HPCs (WT-Ctrl, WT-NMyc, Het-Ctrl, Het-NMyc, Hom-Ctrl, Hom-NMyc). **(B)** Heatmap of differentially expressed genes in the mitochondrial respiratory complex gene set (GO). n=2-3 biologic replicates per group. Groups of genes showing relatively higher levels of expression in Hom-NMyc HPCs are highlighted with the thick black lines.

The pathways in Cluster 1 were upregulated in NMyc-expressing HPCs compared to their controls, often with Hom HPCs being somewhat more “responsive” to NMyc than WT or Het HPCs. This cluster included MYC targets, unfolded protein response genes, as well as genes involved in fatty acid metabolism and oxidative phosphorylation (**Fig. 4A, Fig. S4A–C, and Table S2**). A similar pattern was observed for genes related to the tricarboxylic acid (TCA) cycle (from GO gene sets) (**Fig. S4D and Table S2**). A heat map of differentially expressed nDNA-encoded genes involved in OXPHOS highlighted the more pronounced NMyc-associated upregulation of genes involved in CI and CIV assembly and function in Hom HPCs (**Fig. 4B**). The expression of genes involved in glycolysis was upregulated in Hom-NMyc HPCs but not in WT-NMyc HPCs compared to their respective controls (**Fig. 4A and Fig. S4E),** consistent with the NMyc-induced increase in glycolysis based on the ECARs in Hom but not WT HPCs.

### NMyc expression and the accumulation of somatic mtDNA mutations alter glucose metabolism

We next sought to further understand the link between NMyc-driven metabolic dysregulation and somatic mtDNA mutations in HPCs. Given the significant alterations in the activities and gene expression of aerobic glycolysis and OXPHOS complexes, we employed stable isotope tracing with ^13^C-labled glucose to assess glucose utilization efficiency in ex vivo HPC cultures (**Fig. 5A**). Analysis of ^13^C enrichment in downstream metabolites of glucose oxidation revealed profound differences among NMyc-expressing WT, Het and Hom HPCs. First, we observed that NMyc expression increased the levels of the ^13^C-Ser m+3 isotopologue in WT and Het HPCs, but no such change was detected in Hom HPCs (**Fig. 5A and S5B**). While serine biosynthesis branches from glycolysis, we did not observe changes in other glycolytic intermediates such as 3-phosphoglycerate (3PG) or pyruvate. Second, we observed an increase in both lactate production and glucose oxidation through the TCA cycle in NMyc-expressing WT or Het HPCs compared to their control cells, shown as enhanced ^13^C labeling in the m+3 isotopologue of lactate and the m+2 isotopologues of TCA intermediates (**Fig. 5B, C**). In contrast, NMyc expression failed to increase ^13^C labeling in the m+2 isotopologues of TCA intermediates in Hom HPCs and had a smaller effect on the ^13^C-Lac m+3 isotopologue, which was already increased in Hom-Ctrl HPCs (**Fig. 5C, D**). These results provide additional evidence that the high burden of mtDNA mutation impairs mitochondrial activity in Hom HPCs. Given the critical role of the pyruvate dehydrogenase (PDH) complex, which catalyzes the conversion of pyruvate into acetyl-CoA and therefore links glycolysis to the TCA cycle and OXPOHS within the mitochondria (*47–49*), we further compared PDH activity in NMyc-expressing HPCs by evaluating the Cit m+2/Pyr m+3 ratio. This analysis revealed that NMyc expression substantially increased PDH activity in HPCs of all three genotypes, but to a much lower extent in Hom HPCs (**Fig. 5E**).

**Figure 5.**
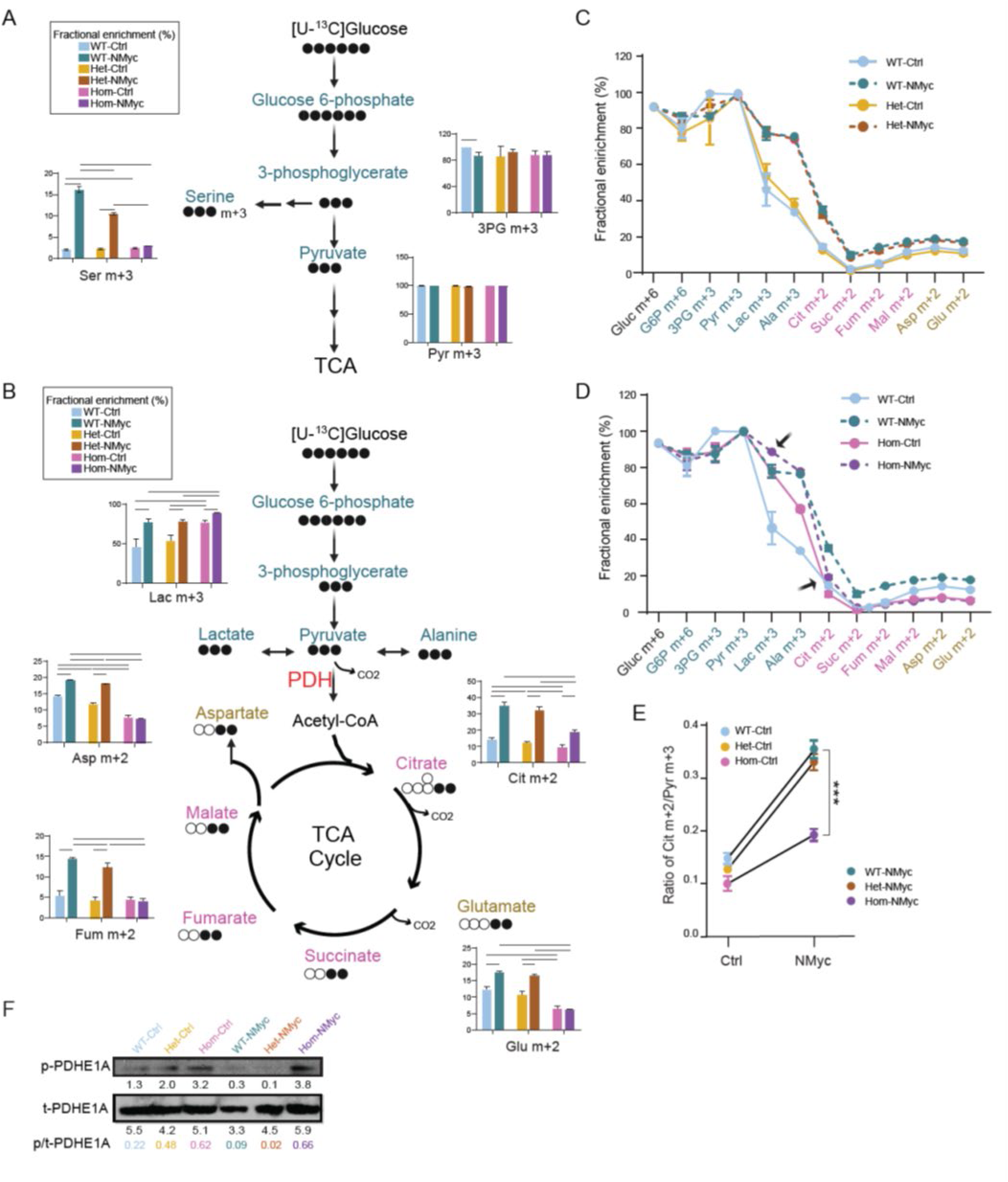
Glucose tracing experiments highlight the resistance of Hom HPCs (but not Het HPCs) to changes in glucose utilization induced by NMyc expression. **(A)** Schematic representation of the glycolytic pathway and its branch to serine biosynthesis with ^13^C-labeled isotopologues. WT, Het, and Hom HPCs (with or without NMyc expression) were incubated with [U-^13^C] glucose, and the incorporation of ^13^C into glycolytic intermediates and serine was analyzed. The schematic highlights the following ^13^C-labeled isotopologues: Glucose (Gluc) m+6, Glucose-6-phosphate (G6P) m+6, Serine (Ser) m+3, 3-phosphoglycerate (3-PG) m+3, and Pyruvate (Pyr) m+3. Bar graphs show fractional enrichment of key ^13^C-labeled isotopologues. n=2-3 biological replicates per group. **(B)** Schematic representation of the glycolytic pathway extending to the tricarboxylic acid (TCA) cycle with ^13^C-labeled isotopologues. WT, Het, and Hom HPCs (with or without NMyc expression) were incubated with [U-^13^C] glucose, and the incorporation of ^13^C into both glycolytic and TCA cycle intermediates was analyzed. The schematic highlights the following ^13^C-labeled isotopologues not shown in **(A)**: Lactate (Lac) m+3, Citrate (Cit) m+2, Succinate (Suc) m+2, Fumarate (Fum) m+2, Malate (Mal) m+2, Aspartate (Asp) m+2, Glutamate (Glu) m+2. Bar graphs show fractional enrichment of key ^13^C-labeled isotopologues. n=2-3 biological replicates per group. The results of Student’s *t-*test are provided in **Table S3. (C, D)** Summary of fractional enrichment across key metabolites, providing a comprehensive overview of ^13^C incorporation in glycolysis, the TCA cycle, and amino acid synthesis. Results for WT-Ctrl, WT-NMyc, Het-Ctrl, and Het-NMyc are shown in **(C)**; results for WT-Ctrl, WT-NMyc, Hom-Ctrl, and Hom-NMyc are shown in **(D). (E)** The ratio of Cit m+2/Pyr m+3 was assessed to compare pyruvate dehydrogenase (PDH) activity in the indicated HPCs transduced with or without NMyc. ***P<0.001. **(F)** Immunoblot analysis using antibodies against phospho-PDHE1A and total-PDHE1A were performed using lysates from sorted YFP+ HPCs. Results of a representative immunoblot analysis and calculated phospho-PDHE1A /total-PDHE1A ratios are shown.

The activity of the PDH complex can be inhibited through phosphorylation of the E1 alpha subunit (p-PDHE1a) at multiple sites by PDH kinases (PDK), which are activated in response to various signals, including the accumulation of NADH (*50, 51*). As NADH levels are often increased in cells with mtDNA mutations affecting CI (*51, 52*), we hypothesized that the impaired activation of PDH in NMyc-expressing Hom HPCs (**Fig. 5E**) might be related to altered PDK activity. To test this hypothesis, we first examined p-PDHE1a levels via immunoblots analyses in Ctrl and NMyc-expressing WT, Het, and Hom HPCs. Although higher levels of p-PDHE1a were observed in Ctrl Het and Hom HPCs compared to WT, p-PDHE1a levels decreased in NMyc-expressing WT and Het HPCs but remained high in Hom-NMyc HPCs (**Fig. 5F, G**). Therefore, NMyc expression overcame PDK’s inhibitory effect and enhanced PDH activity in WT and Het HPCs. However, the impaired OXPHOS in Hom HPCs led to the activation of PDK, resulting in reduced PDH activity and repressed TCA cycle. Additionally, whereas Het HPCs displayed higher metabolic adaptability in response to NMyc-induced oncogenic stress by increasing glucose-derived oxidative flux through the TCA cycle and serine biosynthesis pathway despite initial mtDNA mutation-associated impairments in OXPHOS complex activity, Hom HPCs were unable to do so due to more severe OXPHOS defects associated with a higher burden of mtDNA mutations.

### Impaired NMyc-induced PDH activation limits growth in Hom HPCs

Focusing on WT and Hom HPCs, we next sought to further investigate the contribution of PDK-mediated phosphorylation on PDH activity and its effects on NMyc-induced cellular growth. To this end, we cultured WT and Hom HPCs from both the Ctrl and NMyc groups for 16 h with increasing doses of the PDK inhibitor dichloroacetate (DCA) (*53–55*) in liquid media containing growth factors sufficient to support the BM-derived HPCs **(Fig. 6A and S6A)**. Immunoblot analyses confirmed the higher levels of phosphorylated PDHE1a in NMyc-expressing Hom HPCs than in WT HPCs, as well as the dose-dependent decrease in phosphorylated PDHE1a upon treatment with DCA (**Fig. S6B**). Although the effects of DCA on the phosphorylation of PDHE1a were more pronounced in WT HPCs (with or without NMyc expression) than in Hom HPCs (**Fig. S6B**), NMyc-expressing Hom HPCs were more sensitive to the effects of DCA on cell viability than any of the other HPCs (**Fig. S6C**) when grown in liquid cultures. These results suggest an increased reliance of NMyc-expressing Hom HPCs on glycolysis compared to their WT counterparts.

**Figure 6.**
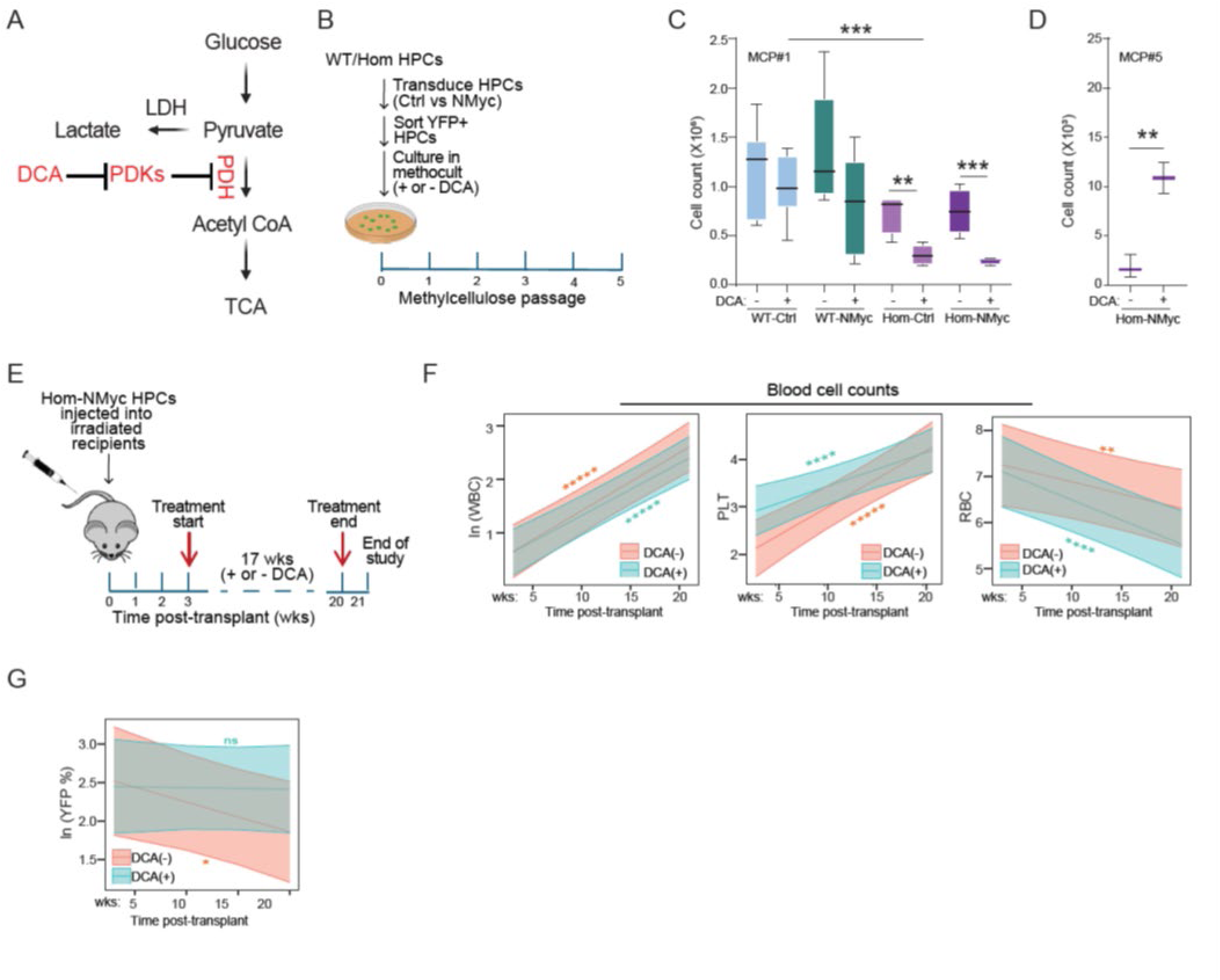
Impaired growth of Hom-NMyc HPCs is partially rescued by inhibiting PDH kinase activity. **(A)** Schematic representation of PDH regulation and the rationale for dichloroacetate (DCA) treatment. Pyruvate derived from glucose is transported into the mitochondria, where it is metabolized by the PDH complex to form acetyl-CoA, which then enters the TCA cycle. PDH activity can be inhibited by factors such as PDH kinases (PDKs), which inactivate the PDH complex by phosphorylating E1α subunit. DCA treatment inhibits PDKs, thereby restoring PDH activity and facilitating the entry of pyruvate into the TCA cycle. **(B)** Schematic of the in vitro experiments to determine the effects of DCA treatment on the growth of HPCs serially passaged in methylcellulose media. Cells were replated in methylcellulose media once a week for 5 weeks (i.e. MCP#1-5). **(C)** The number of HPCs in each treatment group (mean ± s.e.m.) was determined after the first passage (MCP#1), at which time point control and NMyc-expressing Hom HPCs showed exquisite sensitivity to DCA. Sample numbers (biologic replicates): WT-Ctrl (n=6), WT-NMyc (n=6), Hom-Ctrl (n=4), and Hom-NMyc (n=4). All significant differences were measured by ANOVA; **P<0.01, ***P<0.005. **(D)** The number of Hom-NMyc HPCs remaining after MCP#5 (mean ± s.e.m.) was determined, highlighting a reversal in the effect of DCA on the transformation potential of Hom-HPCs. Sample numbers: Hom-Ctrl (n=4) and Hom-NMyc (n=4). The significant difference was measured by ANOVA; **P<0.01. **(E)** Schematic of the experimental design for determining the effects of DCA treatment on the growth of Hom-NMyc HPCs in vivo. HPCs were collected from Hom donors and transduced with NMyc, then subsequentially injected into partially irradiated recipients (802 RAD). Beginning at 3 weeks post-transplantation, recipients were separated into two groups receiving either water containing 5 mM DCA (n=20) or vehicle water (n=15) for 17 weeks. Peripheral blood for flow cytometric and CBC analyses were collected at regular intervals (i.e., 3, 7, 10, 13, 15, 17 and 20 weeks). Sick mice were euthanized upon reaching humane endpoints. The study was terminated at 21 weeks post-transplantation. **(F, G)** Peripheral blood cell counts (WBC, PLT, RBC) **(F)** and the percentage of NMyc expressing (YFP+) cells **(G)** were assessed in two groups of recipients with (n=19) or without (n=13) DCA treatment. Statistical significance was tested using linear mixed-effect models. *P<0.05, **P<0.01, ***P<0.005, ****P<0.001, *****P<0.0005.

To examine the impact of DCA treatment on leukemogenic potential, we seeded HPCs in methylcellulose in the presence or absence of DCA and monitored cell growth upon serial passaging (**Fig. 6C**). The pattern of sensitivity to DCA was different when HPCs were passaged in methylcellulose compared to liquid culture. First, DCA inhibited the colony formation of WT and Hom HPCs, although the effects were more pronounced in Hom HPCs regardless of NMyc expression. More surprising, however, was the reversal of this effect in NMyc-expressing Hom HPCs by the fifth passage in methylcellulose (**Fig. 6D**).

To determine if DCA treatment might have a similar effect on the growth of Hom-NMyc HPCs *in vivo*, we subjected irradiated recipients to DCA treatment beginning 3 weeks after injection with NMyc-expressing Hom HPCs for 17 weeks (**Fig. 6E**). Sub-lethal doses of irradiation were used to condition recipients, avoiding early lethality due to engraftment failure after transplanting recipients with HPCs from Hom mice. Peripheral blood was collected at defined intervals to monitor blood counts (i.e., WBC, PLT, RBC) and assess the percentage of transduced cells (based on the percentage of YFP+ cells) in the recipients. There was no significant difference in survival probability (p=0.23, **Fig. S6D**) during the 21-week study period between recipients treated with DCA and untreated controls. The WBC (and PLT) counts showed similar increases in both groups (**Fig. 6F**); however, the percentage of YFP+ cells in the peripheral blood, which declined significantly in untreated mice (P=0.03) remained stable in those treated with DCA (**Fig. 6G**). Curiously, whereas DCA treatment helped to maintain peripheral levels of transformed Hom-NMyc leukocytes, the decline in RBC counts was more pronounced in those mice (p=0.00002) (**Fig. 6F**). This suggests that although the diversion of glucose into the TCA cycle is beneficial to the process of leukemic transformation, it is detrimental to RBC production in Hom-NMyc mice. Together, the results of the *in vitro* and *in vivo* studies highlight the ability of HPCs to increase glucose-derived oxidative flux through the TCA cycle as a critical component of the metabolic rewiring necessary for NMyc-induced transformation, which is impaired in Hom HPCs.

## DISCUSSION

The metabolic plasticity of tumor cells, which allows them to switch between glycolysis and OXPHOS depending on the context, is crucial for their survival and proliferation (*56–58*). This adaptability ensures that tumor cells can generate sufficient energy and the biosynthetic intermediates required for growth under varying environmental conditions (*59, 60*). Our study highlights the important role of this metabolic flexibility in HPCs and its impact on leukemic transformation driven by oncogenes like NMyc, particularly in the presence of pathogenic mtDNA mutations **(Fig. 7).**

**Figure 7.**
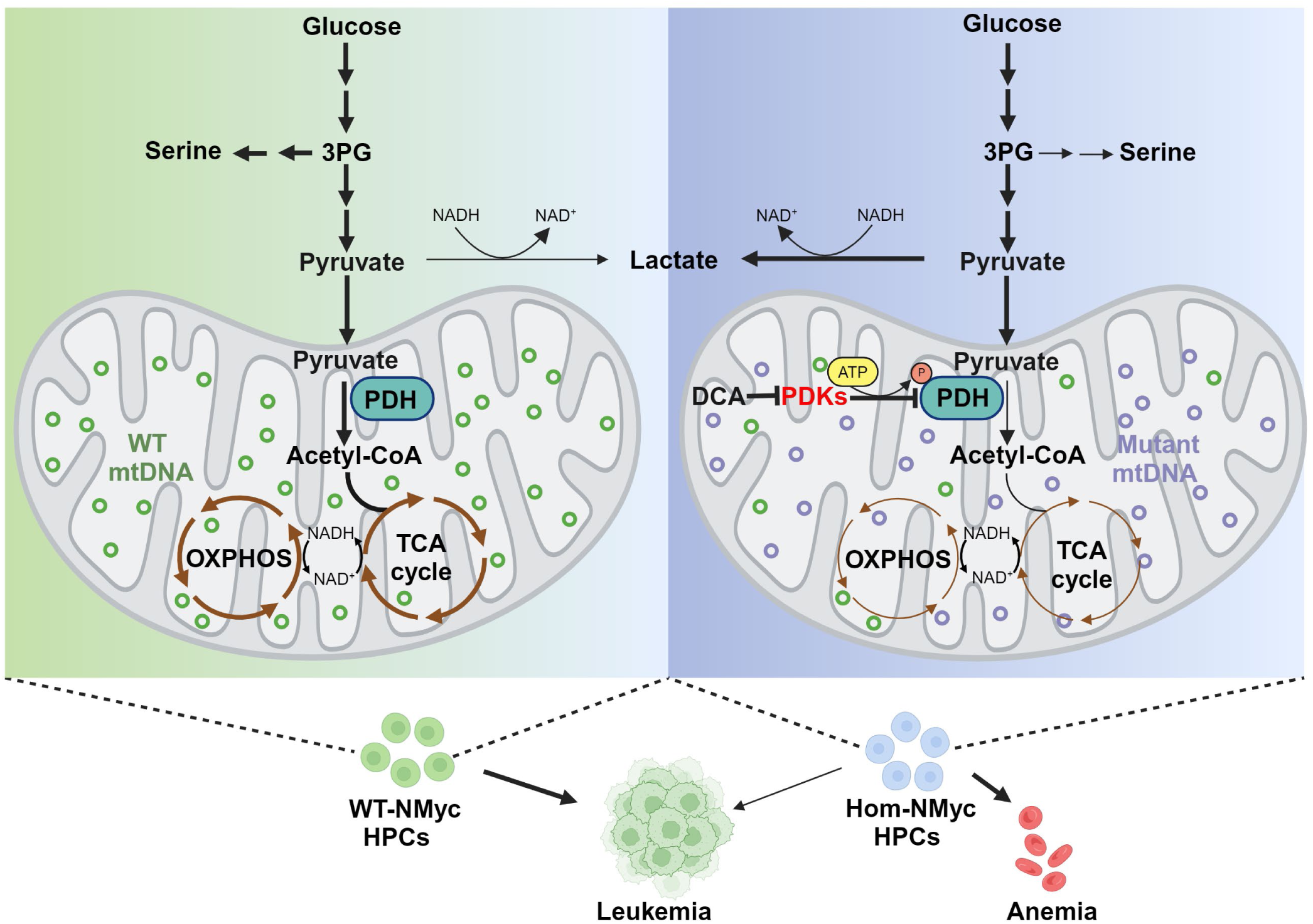
Model summarizing the effects of mtDNA mutation burden on NMyc-induced metabolic rewiring. In WT HPCs (left), NMyc drives the metabolism of glucose towards serine biosynthesis and the production of pyruvate in the cytoplasm through glycolysis. Once transported into the mitochondria, pyruvate is converted to acetyl-CoA by the pyruvate dehydrogenase (PDH) complex generating electron carriers such as NADH (from NAD+). Acetyl-CoA enters the TCA cycle, where it is further oxidized, producing more NADH (and FADH₂) from NAD⁺. Thus, the reaction catalyzed by the PDH complex is pivotal in linking glycolysis with the TCA cycle. The oxidation of NADH (to NAD⁺) is coupled to ATP production during mitochondrial respiration (i.e. OXPHOS). This regeneration of NAD⁺ is necessary for glycolysis and the TCA cycle to continue. NMyc’s ability to promote leukemic transformation NMyc’s ability to promote leukemic transformation depends on the proper interactions between the TCA cycle and oxidative phosphorylation (OXPHOS) in the mitochondria, as these interconnected pathways are crucial for efficient energy production. In NMyc-expressing Hom HPCs (right), pyruvate metabolism is impaired before entering the TCA cycle due to decreased PDH activity, resulting from increased phosphorylation of PDH by pyruvate dehydrogenase kinases (PDKs). The activation of PDK in Hom HPCs, which is further exacerbated by NMYC, is likely related to the accumulation of NADH that occurs with disruption of mitochondrial respiration (e.g. due to the heavy burden of mtDNA mutations). The PDK-mediated inhibition of PDH complex activity decreases the flux of glucose-derived metabolites into the TCA cycle. As a result, pyruvate is preferentially converted to lactate via lactate dehydrogenase (LDH) to regenerate NAD⁺ from NADH in the cytoplasm. This conversion is crucial because it maintains the NAD⁺/NADH ratio, allowing glycolysis to continue despite the impaired mitochondrial function. The impaired serine biosynthesis may also be related to disruptions in the NAD+/NADH ratio and/or the shift towards glycolysis. In Het HPCs (not shown), exposure to a low level of mtDNA mutations might confer a degree of metabolic flexibility or plasticity. These cells can potentially adapt to minor mitochondrial dysfunctions, maintaining a balance between glycolysis and the TCA cycle. This metabolic adaptability may provide a survival advantage under varying metabolic conditions, allowing Het HPCs to better support leukemic transformation compared to Hom HPCs, which are overwhelmed by the high mutation burden.

We found that the burden of somatic mtDNA mutations influences the leukemogenic potential of NMyc in HPCs. Het HPCs, with an intermediate burden of mtDNA mutations, showed minimal impairments in supporting NMyc-dependent transformation. In contrast, Hom HPCs, with a higher burden of mtDNA mutations, exhibited a marked reduction in their leukemogenic potential. Moreover, Het HPCs demonstrated a higher frequency of spontaneous leukemic transformation compared to WT cells, suggesting that an optimal level of mitochondrial dysfunction may create a conducive metabolic environment for oncogenic transformation, whereas excessive mitochondrial dysfunction is detrimental (*61*).

Our findings show both similarities and differences with previous studies that used the mutant *Polg* expressing mtDNA mutator mice to modulate mtDNA mutation burden. For instance, the cancer predisposition phenotype in a Li-Fraumeni syndrome mouse model with mutant p53 is reduced in Hom *Polg* mutant mice (*62*), indicating that excessive mtDNA mutations can impair tumorigenesis. Nevertheless, in this system, tumorigenesis was also impaired (albeit to a lesser degree) in Het *Polg* mutant mice, suggesting a dose-dependent effect of mtDNA mutations on transformation potential –-compared to the threshold effect of mtDNA mutation burden on leukemic transformation in our model. Moreover, the increased susceptibility to colon cancer in an inducible intestinal tumorigenesis model observed in Hom *Polg* mice (*63*), and the increase in spontaneous leukemogenesis that we observed using HPCs from Het *Polg* mice, suggest that a certain level of mitochondrial dysfunction can promote transformation in specific contexts (*19, 20*). These observations underscore the complex and context-dependent role of mtDNA mutations in cancer development, highlighting the importance of balancing mitochondrial function and dysfunction in determining oncogenic outcomes.

Our *in vitro* studies further characterize the relationship between mtDNA mutations and leukemic transformation. We observed that the ability of HPCs to support NMyc-driven transformation correlates with their capacity to overcome the effects of mtDNA mutations on OXPHOS and glucose metabolism. While both Het- and Hom-Ctrl HPCs exhibited comparable decreases in CIV activity, only Het HPCs managed to enhance CIV activity to meet the increased oxidative metabolism demands imposed by NMyc. Het HPCs also increased glucose-derived oxidative flux through the TCA cycle and serine biosynthesis pathway in response to NMyc expression, achieving levels comparable to those in NMyc-expressing WT HPCs. In contrast, Hom HPCs favored glycolysis and failed to adapt to the metabolic demands of NMyc expression.

We speculate that dysregulation of the NAD+/NADH ratio is a critical factory underlying the impaired metabolic adaptability of Hom HPCs (**Fig. 7**). Such dysregulation could account for the impaired conversion of 3-phosphoglycerate to serine. Phosphoglycerate dehydrogenase (PHGDH), which catalyzes the first and rate-limiting step in the serine biosynthesis pathway by converting 3-phosphoglycerate to 3-phosphohydroxypyruvate, relies on NAD+ as a cofactor and is inhibited when NADH levels are elevated (*64*). Concurrently, PDK, which inhibits oxidative glucose flux through the TCA cycle, is activated under increased NADH levels (*65*). Therefore, it is plausible that the altered NAD+/NADH ratio associated with impaired OXPHOS in Hom-NMyc HPCs compromises their metabolic plasticity and ability to undergo leukemic transformation.

Interestingly, leukemia derived from Hom HPCs lacks mutations in critical oncogenic pathways such as PTEN/PI3K/AKT, RAS, and p53, which are commonly mutated in acute leukemias and promote cell survival, proliferation, and metabolism (*36–39*). This suggests that the high burden of mtDNA mutations in Hom HPCs imposes significant metabolic constraints, preventing the acquisition of additional mutations within these pathways. The compromised OXPHOS capacity and unfavorable NAD+/NADH ratio in Hom HPCs likely disrupt key metabolic and biosynthetic pathways, creating a metabolic bottleneck that limits their oncogenic adaptability and potential for malignant transformation.

Conversely, the increased incidence of spontaneous leukemia in Het HPCs suggests that their metabolic plasticity not only exceeds that of Hom HPCs but also surpasses that of WT HPCs. The intermediate burden of mtDNA mutations in Het HPCs may foster a unique metabolic environment that supports both glycolytic and oxidative phosphorylation pathways, enhancing their ability to switch between metabolic states to meet varying energetic and biosynthetic demands (*66–68*).This enhanced metabolic plasticity could provide a selective advantage under oncogenic stress, making Het HPCs more permissive to transformation by a broader range of oncogenes. Our observations align with this hypothesis, as Het HPCs support NMyc-driven transformation more effectively than Hom HPCs and exhibit a higher frequency of spontaneous leukemic transformation compared to WT HPCs.

This study underscores the critical role of metabolic adaptability in leukemogenesis and highlights the complex interplay between mtDNA mutations and oncogenic transformation. The intermediate mtDNA mutation burden in Het HPCs primes these cells for oncogenic transformation, while excessive mtDNA mutations in Hom HPCs impose metabolic constraints that limit their leukemogenic potential. Further research is warranted to elucidate the specific metabolic pathways and oncogenic signals preferentially exploited in Het HPCs, which could inform novel therapeutic strategies targeting metabolic vulnerabilities in leukemia (*68*).

## MATERIALS AND METHODS

### Mice

*PolgA^D257A^*-heterozygous mice (*Polg^wt/mt^*) (*21*) were maintained by backcrossing to C57BL/6J (JAX Stock# 000664, The Jackson Laboratory, Bar Harbor, ME, USA) and interbred to generate *Polg^mt/mt^*(Hom), *Polg^wt/mt^* (Het), and *Polg^wt/wt^* (WT) littermates that were used in experiments. Recipients C57/B6 mice (JAX Stock# 000664) were purchased from The Jackson Laboratory. Cohorts of 5–15 recipients per donor group (i.e., WT, Het, or Hom HPCs with (NMyc) or without (control; Ctrl) ectopic NMyc expression) from 2- to 5-month-old mice were generated from at least five independent experiments and were designated as follows: WT-Ctrl (n = 39), Het-Ctrl (n = 36), Hom-Ctrl (n = 27), WT-NMyc (n = 82), Het-NMyc (n = 83), and Hom-NMyc (n = 89) (Fig. 1B). Mice were housed in a 12-hour light-dark cycle and were provided food and water *ad libitum*. All animal experiments were approved by the Institutional Animal Care and Use Committee (IACUC# 601-100574) at St. Jude Children’s Research Hospital (SJCRH).

### Cell sorting

Lineage-negative BM cells (hereafter referred to as HPCs) were separated by the autoMACS^®^ Pro Separator (Miltenyi Biotec, Bergisch Gladbach, Germany), with depletion of lineage-positive cells using anti-biotin–conjugated magnetic beads (130-090-485, Miltenyi Biotec) and biotin-labeled mouse antibodies (BD Biosciences (Franklin Lakes, NJ, USA) or BioLegend (San Diego, CA, USA)) against CD4, CD8, NK1.1, B220, Mac1, Gr1, and Ter119. All sorted HPCs were either immediately seeded for 48 h in complete BM (c-BM) media (Iscove’s Modified Dulbecco’s Medium with 20% fetal calf serum, 50 ng/mL murine stem cell factor, 25 ng/mL murine interleukin-3, and 25 ng/mL murine IL-6) for further retroviral transduction or kept on ice for cell pellet preparation.

### Vector constructs and retroviral transduction

Hemagglutinin (HA)-tagged WT *NMyc* cDNA(*25*) was cloned into the *Eco*RI site of the mouse stem cell virus–internal ribosome entry site–yellow fluorescent protein (MSCV-IRES-YFP) retroviral vector. HA-tagged mutant NMyc cDNAs lacking sequences encoding the NH_2_-terminal transactivation/repression were generated via PCR and cloned into the same *Eco*RI site 5′ of the IRES sequence of MSCV-IRES-YFP, thereby generating MSCV-NMyc-IRES-YFP.

GP+E86 cells expressing the MSCV-IRES-YFP or MSCV-NMyc-IRES-YFP construct were kindly gifted by Dr. John Schutz (SJCRH). The GP+E86 cells with MSCV-IRES-YFP or MSCV-NMyc-IRES-YFP were maintained in culture for at least 2 days prior to spin transduction. Six-well plates were coated with retronectin (50 µg/2 mL/well, (Takara Bio Inc., Kusatsu, Japan)) for 16 h, followed by spin transduction with supernatant containing MSCV-IRES-YFP (Ctrl) or MSCV-NMyc-IRES-YFP (NMyc) at 3000 rpm, 60 min, 4°C. The supernatants from these cells were used to transduce HPCs as previously described by (*40*). The HPCs from WT, Het, and Hom mice were then seeded on the plates, with 0.5 × 10^6^ HPCs/well in c-BM media. Transduced HPCs from both groups were injected into lethally irradiated recipients; additional YFP+ cells were sorted for further *in vitro* assays, such as colony-forming cell assays (see below) (Fig. 1A).

### Bone Marrow Transplantation

Retrovirally transduced HPCs cells (1 × 10^6^) were intravenously injected into lethally irradiated (9.8 Gy) syngeneic recipients. Recipients were closely monitored post-transplantation and sacrificed upon reaching humane endpoints or at 250 days post-transplantation, whichever occurred earlier. At that time, peripheral blood was collected for flow cytometric analyses and complete blood counts (see Fig. S1D–G) and full necropsies were performed. A veterinary pathologist blinded to the experimental design assessed histological samples to determine the extent of leukemic infiltration and reported any other pathological findings.

### Colony-forming assay

After sorting, the YFP HPCs were grown in c-BM media for 72 h, 1 × 10^3^ cells were suspended in 100 µL c-BM media with 1 mL MetholCult^TM^ GF M3434 media (Stemcell Technologies, Vancouver, BC, Canada) and then maintained in culture on a 35-mm plate (Cat# 27150, Stemcell Technologies) in triplicate. Colonies and cell numbers were scored for each plate after 7 days. Colonies containing at least 10 cells were scored. Serial replating assays were performed, in which the cells were harvested and replated every 7 days until week 6 of methylcellulose assays (MCP#1 to MCP#6).

### Tissue collection

Mice were euthanized with CO_2_, followed by cervical dislocation. Peripheral blood was collected via retro-orbital bleeding, and complete blood counts were analyzed by the Veterinary Pathology Laboratory Core at SJCRH. BM or spleens were harvested and stored on ice for immediate use or were frozen for later experiments. Single-cell suspensions were prepared for flow cytometry and cell-sorting experiments. Frozen BM cell pellets and snap-frozen tissues were stored at −80°C until needed for extraction of DNA, RNA, and protein.

### Histopathology and immunohistochemistry

The liver, spleen, sternum, and any tumor tissues were sampled and postfixed in 10% formalin and before being sent to the Veterinary Pathology Core at SJCRH for histologic analyses. Briefly, all tissues were fixed in formalin, embedded in paraffin, sectioned at a thickness of 4 μm, mounted on positively charged glass slides (Superfrost Plus; Thermo Fisher Scientific, Waltham, MA, USA), and dried at 60°C for 20 min before dewaxing and staining with hematoxylin and eosin using standard methods. For immunohistochemical staining, the following primary antibodies were used: for T cells, anti-CD3 (1:1000 dilution, Cat # sc-1127; Santa Cruz Biotechnology (Dallas, TX, USA)); for B cells, anti-B220 (1:1,000 dilution, cat# 553084; BD Biosciences) and anti-PAX5 (1:1000 dilution, Cat # ab109443; Abcam (Cambridge, UK)); for myeloid cells, anti-myeloperoxidase (1:1200 dilution, cat# A0398; Agilent (Santa Clara, CA, USA)); for N-myc detection, anti-HA (1:1000 dilution, C29F4, # 3724; Cell Signaling Technology (Danvers, MA, USA)). Tissue sections underwent antigen retrieval in a prediluted cell-conditioning solution (*CC1;* Ventana Medical Systems (Oro Valley, AZ, USA)) for 32 min. The OmniMap anti-Rabbit HRP kit (Ventana Medical Systems) and ChromoMap DAB (Ventana Medical Systems) were used for detection. All sections were examined by a pathologist blinded to the experimental group assignments.

### Flow cytometry

Red blood cells (RBCs) were lysed and eliminated from peripheral blood samples using RBC lysis buffer (Cat: 420301, BioLegend). Expression of YFP and cell-surface markers was determined by flow cytometry with the following antibodies: CD3e (145-2C11 APC, BD 553066), B220 (CD45R, RA3-6B2 Brilliant Violet 605, BioLegend 103244), CD4 (L3T4, RM4-5 PE, BD 553049), CD8a (Ly2, 53-6.7 PE-Cy7, BD 552877), Sca-1 (Ly-6 A/E, D7 PerCP-Cy5.5, eBioscience 45-5981-82, 100 ug), CD11b (Mac-1, M1/70 V500, BD 562127), Gr1 (RB6-8C5, Alexa 700, BD 557979), CD19 (clone 1D3, BV650, BD 563235), and CD117 (c-Kit, 2B8 APC-eFluor 780, eBioscience 47-1171-82). All fluorescence-activated cell sorting data were analyzed using FlowJo software (BD Biosciences).

### RNA and gene expression profiling

Total RNA (100 ng) was isolated from HPCs using the RNeasy Mini Kit (QIAGEN, Hilden, Germany). Sense target cDNA was prepared using the Affymetrix GeneChip WT Plus kit (Applied Biosystems P/N 902281). It was then fragmented and biotin-labeled using the GeneChip WT Terminal Labeling kit (Applied Biosystems P/N 900670) and then hybridized to a Clariom S Mouse GeneChip array (Applied Biosystems P/N 902931). Probe signals were normalized and summarized to log_2_-expression values by using a robust microarray analysis (*69*) and the Affymetrix Expression Console software v1.1. Initial exploration of the expression data by principal component analysis identified appreciable variation associated with experiment date. Therefore, the experiment date was included in an ANOVA designed to identify profiles that varied according to: (1) *Polg* genotype, (2) NMyc expression, and (3) profiles that depend on the interaction between *Polg* and NMyc expression. A three-factor ANOVA (genotype, treatment, experiment date) was used to perform gene set analysis, as implemented within Partek Genomics Suite v7.0 (Partek, Inc., San Diego, CA, USA). Gene sets were derived from GO categories, C2-Canonical pathways, and Hallmark pathways. Figures and supplemental tables show the mean expression values and differences, as calculated by an ANOVA. A false discovery rate of less than 5% was applied to report significant findings.

### gDNA extraction

BM cells collected at different stages (primary HPCs, MCP#0, and total BM cells at from sick mice) were used for genomic DNA (gDNA) extraction by using the Qiagen kit (DNeasy Blood & Tissue, cat# 69504) per the manufacturer’s instructions. All gDNA samples were quantified using a Thermo Fisher Scientific Nanodrop 1000 (Thermo Fisher Scientific) and were stored at –20°C until further use.

### PCR amplification of mtDNA

Following the gDNA extraction, long-range PCR (LR-PCR) amplification was performed with two primers against mtDNA (Fw: ctacctttgcacggtcaggatacc 2033-2010, Rev: cggctaaacgagg gtccaactgt c 2071-2094, region 2094–2033) to amplify 16-kb amplicons by using TaKaRa LA Taq® DNA Polymerase Hot-Start Version (Takara Bio Inc., RR042A) (*30*). The PCR reaction was executed using 50-μL mixtures, including 5 µL 10× LA PCR buffer, 8 µL dNTP mixture, 1 µL 10 pM primer pairs, 0.5 µL TaKaRa LA *Taq HS*, and 10 ng genomic template DNA. Thermal cycling conditions included an initial denaturation at 95°C for 2 min, then 30 cycles at 95°C for 20 s, followed by 68°C for 18 min, and a final extension at 68°C for 20 min. PCR amplicons were quantified using a Thermo Fisher Scientific Nanodrop 1000 (Thermo Fisher Scientific) and sent to the Hartwell Center for Biotechnology at SJCRH for Mito-Seq.

### Mitochondrial DNA Sequencing and analysis

Amplicons were purified using an 0.8× Ampure XP cleanup following the manufacturer’s instructions (Beckman Coulter Life Sciences, Brea, CA. USA). Libraries were prepared from 1 ng amplicon DNA by using the Nextera XT library prep kit (Illumina, San Diego, CA, USA) according to the manufacturer’s instructions. Libraries were analyzed for insert-size distribution on a 2100 BioAnalyzer High Sensitivity kit (Agilent), a 4200 TapeStation D1000 ScreenTape assay (Agilent), or a Caliper LabChip GX DNA High Sensitivity Reagent Kit (PerkinElmer, Waltham, MA, USA). Libraries were quantified using the Quant-iT PicoGreen ds DNA assay (Life Technologies, Carlsbad, CA, USA). Paired-end, 150-cycle sequencing was run on a MiSeq using a v2 Micro kit (Illumina).

After removing the adapter sequences and trimming low-quality bases, sequencing reads (Phred-like score Q25 or greater) with at least 50 nucleotides of whole exome sequencing data from 41 mice (12 WT, 12 Het, 6 Hom, and 11 donor controls) were mapped to mouse genome NCBI37/mm9 reference to achieve an average depth coverage above 150× for the targeted exon regions by using CLC Genomics Workbench v11.0 (CLCbio, Qiagen). The single nucleotide variants and short indels were then called using the Variant Detection tool. After filtering against samples of controls, mutations with at least 10% variant allele frequency and 30× depth coverage were summarized in the final variant report. The genes that were mutated with the highest frequency among the 30 experimental mice are highlighted in the list. The significance of mutations involved in certain pathways across genotype groups was determined via Fisher’s exact test.

For 69 mtDNA samples prepared with 150-bp paired-end reads, image analyses and base calling were performed using MiSeq control software and real time analysis. After trimming, high-quality reads with at least 50 nucleotides were aligned to a mouse mtDNA reference sequence. With average sequencing depth of above 1500×, the frequencies of mutation at each sequence position were determined using CLC Workbench. The numbers of mutations in each sample with a variant allele frequency greater than 1%, 5%, or 10% were summarized, and the significant differences in mitochondrial mutation accumulation across samples from different genotypes were determined using ANOVAs.

### Exome sequencing

The gDNA was prepared as described above. Mouse genomic libraries were generated using the SureSelect^XT^ kit specific for the Illumina HiSeq instrument (Catalog No. G9611B; Agilent), followed by exome enrichment using the SureSelect^XT^ Mouse All Exon bait set (Catalog No. 5190-4642). The resulting exome-enriched libraries were then sequenced by the Genome Sequencing Facility at SJCRH. Paired-end 100-cycle sequencing was performed on a HiSeq 2500 or HiSeq 4000 (Illumina), according to the manufacturer’s directions.

### mtDNA quantification

The gDNA samples were diluted in water, and total of 10 ng DNA was analyzed by quantitative PCR using FastStart Universal SYBR Green master mix (Roche, Basel, Switzerland) with primers directed against gDNA or mtDNA. The following primers were used: COI (mitochondrial) (Fw: 5′-GCCCCAGATATAGCATTCCC-3′, Rev: 5′-TTCATCCTGTTCCTGCTCC-3′); ND2 (mitochondrial) (Fw: 5′-CCATTCCACTTCTGATTACC-3′, Rev: 5′-ATGATAGTAGAGTTGAGTAGCG-3′); and 18S (nuclear) (Fw: 5′-TAGAGGGACAAGTGGCGTTC-3′, Rev: 5′-CGCTGAGCCAGTCAGTGT-3′). The mtDNA value was normalized to the 18S RNA (nDNA) value.

### OCRs and ECARs

OCRs and ECARs were quantified using an XFe-24 Extracellular Flux Analyzer (Seahorse Bioscience). Briefly, 1 × 10^5^ cells/well were seeded with XF media (nonbuffered DMEM containing 25 mM glucose, 2 mM L-glutamine, and 1 mM sodium pyruvate) in a Cell-Tak (BD Biosciences)–coated plate 17 h after infection and 1 h before loading the plate into the instrument.

### Citrate Synthase assay

CS activity was assayed in whole-cell lysates described previously (*70*). Samples were diluted in the reaction buffer consisting of 50 mM Tris (pH 8.0), 0.5 mM oxaloacetate, and 0.1 mM 5,5ʹ dithiobis-(2-nitrobenzoic acid). Reactions were performed at 25°C and were initiated by the introduction of 50 mM acetyl-CoA. Absorbance was monitored at 412 nm for 1–2 min by using a Shimadzu UV1800 dual-beam spectrophotometer, and enzyme activity was calculated using the extinction coefficient Ɛ=13.6 × mM^-1^cm^-1^. To ensure reproducibility, activity was measured using a range of protein concentrations for each lysate, and averages were calculated for those with linear specific activity results.

### CIV assay

CIV was assayed in whole-cell lysates by measuring the change in absorbance of ferrocytochrome c. Reaction mixtures consisting of 50 mM Tris (7.5) and 25 µM ferrocytochrome c, which were prepared by reducing a ferricytochrome c stock solution with dithiothreitol, were allowed to equilibrate before use. Reactions were performed at 25°C and were initiated by adding lysate. The absorbance at 550 nm was then measured for 1 min using a Shimadzu UV1800 spectrophotometer. Activity was calculated using the extinction coefficient Ɛ=21.8 × mM^−1^cm^−1^. To ensure reproducibility, activity was measured using a range of protein concentrations for each lysate, and averages were calculated for those with linear specific activity results.

### Immunoblot analyses

Protein extracts were prepared as previously described (*35*). Cleared lysates from cells or tissues were separated on 4%–12% Bis-Tris gels (Invitrogen) and transferred onto a nitrocellulose membrane. After blocking in 5% skim milk or 5% bovine serum albumen, blots were probed using the following primary antibodies per the manufacturer’s instructions: GAPDH (G9545), phosphor-PDHE1-a [p ser293] (NB11093479), anti-PDHA1[8D10E6] (ab110334), LDHa (CS2012), Mito Total Oxphos (ab110413), and Tom20 (FL145, SC11415) (Santa Cruz Biotechnology). Bands were detected by horseradish peroxidase–labeled secondary antibodies and an enhanced chemiluminescence detection kit (Cytiva, Marlborough, MA, USA).

### Metabolomics assays

The protocol was kindly provided by Dr. Daniel Brass (UCLA Metabolomics Center), with minor modifications. Briefly, HPCs were plated in 6-well plates (1 × 10^6^ cells/well; 3 wells per condition) and incubated overnight. Cells were incubated in medium with dialyzed FBS containing labeled glucose 4.5 g/L U-13C6 D-glucose, Cambridge Isotope Laboratories, Inc. (Tewksbury, MA, USA) for 18 h. For cell samples, cells were rinsed with cold 150 mM ammonium acetate. The cells were then scraped off with 1 mL 80% cold methanol. An internal standard of 10 nmol norvaline was added, and the cell suspension was spun down at top speed (12X1000 rpm) for 5 min at 4°C. The supernatant was transferred into a glass vial, and the pellet was resuspended in 200 μL 80% methanol and spun again. The supernatant was then added to the glass vial. The metabolites were dried in an EZ-2Elite evaporator at 30°C using an aqueous program and stored at –80°C until mass spectrometry was performed by the UCLA Metabolomics Center.

Dried metabolites were then resuspended in 50% Acetonitrile, and a one-tenth aliquot was loaded onto a Luna 3-µm NH2 100A (150 × 2.0 mm) column (Phenomenex, Torrance, CA, USA). The chromatographic separation was performed on an UltiMate 3000 RSLC (Thermo Fisher Scientific) with mobile phases A (5 mM NH_4_AcO; pH 9.9) and B (ACN) and a flow rate of 200 μL/min. The gradient from 15% A to 95% A over 18 min was followed by 9-min isocratic flow at 95% A and re-equilibration to 15% A. Metabolite detection was achieved with a Thermo Fisher Scientific Q Exactive mass spectrometer run with polarity switching (+3.0 kV/−2.25 kV) in full-scan mode, with an m/z range of 65-975. TraceFinder 4.1 (Thermo Fisher Scientific) was used to quantify metabolites by area under the curve using expected retention time and accurate mass measurements (<3 ppm). Relative amounts of metabolites were calculated by summing all isotopologues of a given metabolite and normalized to cell number. Data analyses, including principal component analysis and hierarchical clustering, were performed using in-house scripts in the R programming language.

### DCA treatment

DCA (347795_10G, Millipore Sigma) was diluted in different concentrations *in vitro*: 0, 5, or 25 µm for liquid culture and 10 µm for methylcellulose media. *In vivo,* recipients were provided water treatment with supplement 750 mg/L (5 mM) of DCA for a cage of five mice at post-transplantation 14 days. DCA was administered to 20 mice injected with Hom-NMyc HPCs while vehicle water was given to another 15 Hom-Ctrl mice (Fig. 5F). The treatment lasted for 17 weeks (*71*).

### Statistical analyses

Statistical analyses were performed using Prism GraphPad v10 (GraphPad, La Jolla, CA, USA). Significance was assessed using a Student’s *t-*test or one-factor ANOVA (\**p* <0.05), unless otherwise indicated in specific assays. A log-rank test was conducted to examine the difference in survival distributions among the genetic groups (*72*). Multiple Gray’s tests were performed to examine the difference in cumulative incidence rates between the pre-specified comparisons under competing risks (*73*). The cumulative incidence rate was computed for tumor death (event of interest) and other causes (competing risks) in different conditions. Linear mixed-effects models, which handle the correlation of repeated measurements over time, were used to exam the association between YFP, WBC, RBC, PLT and treatment (*74*).

## ACKNOWLEDGMENTS

We thank Daniel Brass (Arrowhead Pharmaceuticals, Madison, Wisconsin) and Joanna Scott (Metabolomics Center, UCLA) for performing the metabolomics assay; Granger Ridout and Scott Olsen (Hartwell Center, St. Jude) for technical support on RNA expression profiling and mitochondrial sequencing; the following shared resources at St. Jude Children’s Research Hospital for experimental assistance: the Cell and Tissue Imaging Center, the Flow Cytometry and Cell Sorting Shared Resource, the Animal Resource Center, the Immunology Flow Cytometry Core, and the Hartwell Center for Bioinformatics and Biotechnology. We thank Danielle D’Amore and Angela McArthur for editing the manuscript. The authors declare no competing interests. All data needed to evaluate the conclusions in the paper are present in the paper and/or the Supplementary Materials. **Funding:** This project received funding from the National Institutes of Health (R01MH115058 and R01GM132231 to M.K., and R01CA194206 to J.S.) and the American Lebanese Syrian Associated Charities (to M.K, M.N. and J.S.)

## AUTHOR CONTRIBUTIONS

- Conceptualization: M.K., and X.L.-H.
- Methodology: X.L.-H., J.L., Y.F., G.P., M.K., H.R., C.W.W., and A.S.
- Investigation: H.W., Y.-D.W., G.N., A.S., S.P., J.S., and M.N.
- Supervision: M.K.
- Writing—original draft: X.L.-H., and M.K.
- Writing—review and editing: X.L.-H., M.N., and M.K.

## SUPPLEMENTAL FIGURES

**Figure S1.**
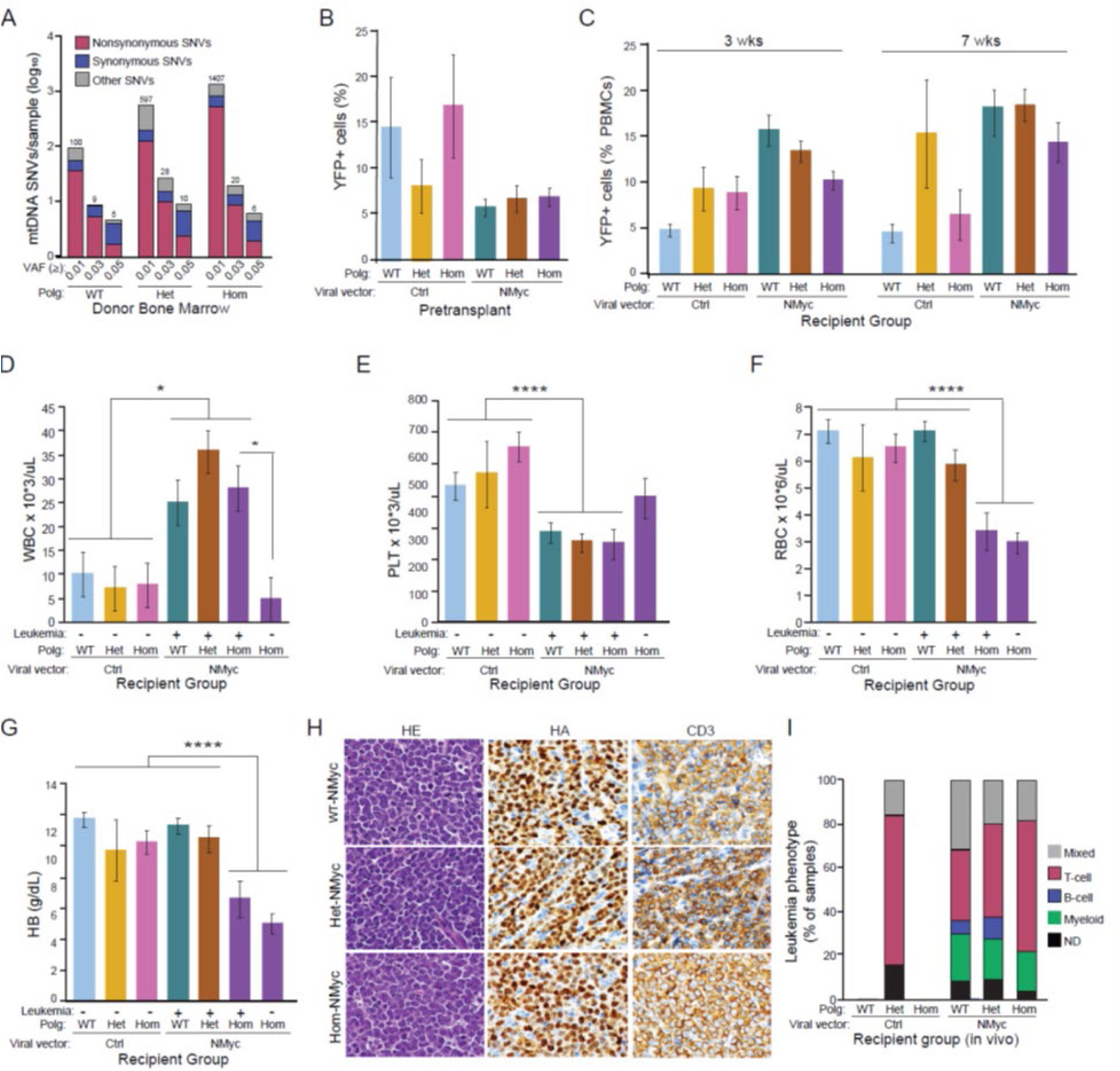
Supplement to Figure 1. **(A)** Bar graph illustrating the frequency of mitochondrial DNA (mtDNA) single nucleotide variants (SNVs) per sample at or below different variant allele frequency (VAF) cutoffs (0.01, 0.03, 0.05). The SNVs are categorized into three classes: synonymous, non-synonymous, and other. The data is derived from bone marrow cells isolated from wild-type (WT), heterozygous (Het), and homozygous (Hom) mice: WT (n=4), Het (n=2), Hom (n=3). The different VAF cutoffs are represented on the x-axis, while the frequency of SNVs per sample is depicted on the y-axis. Each bar is color-coded to distinguish between synonymous, non-synonymous, and other types of SNVs, providing a visual comparison of their relative frequencies across the specified VAF thresholds. **(B)** The percentage of YFP+ hematopoietic progenitor cells (HPCs) was measured using flow cytometry after transduction with either MSCV-IRES-YFP (control; Ctrl) or MSCV-IRES-NMyc-YFP (NMyc). The graph shows the percentage of transduced (YFP+) cells in each group: WT-Ctrl (n=5), Het-Ctrl (n=3), Hom-Ctrl (n=5), WT-NMyc (n=5), Het-NMyc (n=4), and Hom-NMyc (n=5). **(C)** The percentage of YFP+ cells among viable mononuclear cells was analyzed via flow cytometry using peripheral blood samples from different recipients at approximately 3 weeks and 7 weeks post-transplantation. Sample sizes analyzed at 3 weeks post-transplantation were: WT-Ctrl (n=24), Het-Ctrl (n=14), Hom-Ctrl (n=20), WT-NMyc (n=54), Het-NMyc (n=60), and Hom-NMyc (n=58). Sample sizes at 7 weeks post-transplantation were: WT-Ctrl (n=21), Het-Ctrl (n=10), Hom-Ctrl (n=16), WT-NMyc (n=56), Het-NMyc (n=56), and Hom-NMyc (n=47). **(D–G)** Complete blood count (CBC) results for recipients who became sick compared to control mice. Bar graphs showing white blood cell (WBC) count **(D)**, platelet (PLT) count **(E)**, red blood cell (RBC) count **(F)**, and hemoglobin (HB) level **(G)**. Sample sizes were: WT-Ctrl (n=8), Het-Ctrl (n=6), Hom-Ctrl (n=12), WT-NMyc (n=28), Het-NMyc (n=20), Hom-NMyc (with leukemia) (n=8), and Hom-NMyc (without leukemia) (n=29). *P<0.05 and ****P<0.00001 (ANOVA). **(H-I)** Representative images of spleen sections from WT-NMyc, Het-NMyc, and Hom-NMyc mice that reached humane endpoints, stained with hematoxylin and eosin (HE) or antibodies against the T-cell marker CD3 and hemagglutinin (HA) to detect HA-tagged NMyc **(H)**. Similar experiments were conducted on tissues harvested at necropsy to characterize leukemic infiltrates in other tissues, such as bone marrow (BM) and liver, using these markers and additional ones, including B-cell markers (B220 and PAX5) and the myeloid cell marker MPO. Results from these immunohistochemical analyses, along with immunophenotyping of peripheral leukemic blasts (when available) by flow cytometric analysis, were used to classify the leukemic phenotype into one of five categories: T-cell, B-cell, mixed lineage, myeloid, or not determined (ND). The bar graph **(I)** shows the distribution of samples across the phenotypic categories among mice with leukemia in the different groups. Sample sizes related to the incidence of leukemia in the different groups were: WT-Ctrl (n=1), Het-Ctrl (n=5), Hom-Ctrl (n=1), WT-NMyc (n=68), Het-NMyc (n=61), Hom-NMyc (n=14).

**Figure S2.**
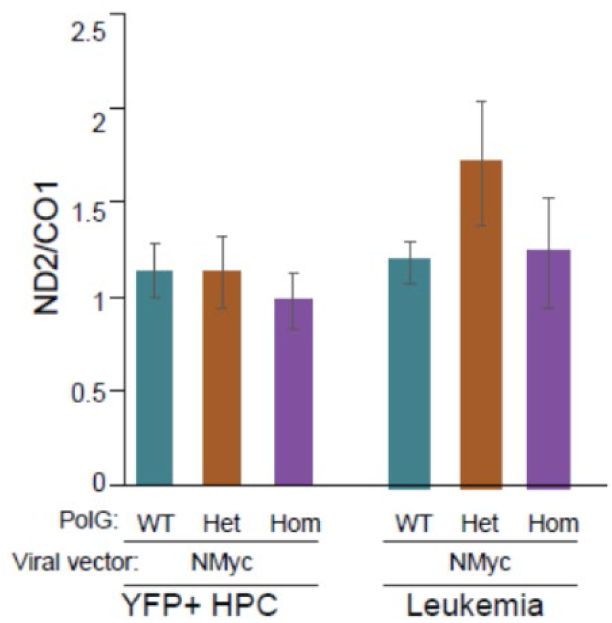
Supplement to Figure 2. Quantification of mtDNA content in YFP+ HPCs prior to transplantation and in cells from leukemia-infiltrated BM samples. The graph shows relative mtDNA copy number (mean ± s.e.m) calculated using ND2 and CO1 as mtDNA markers and 18S and the nDNA marker. The ND2/CO1 ratios were similar across all samples: YFP+ HPCs prior to transplantation - WT-NMyc (n=7), Het-NMyc (n=6), and Hom-NMyc (n=7); leukemic BM - WT-NMyc (n=15), Het-NMyc (n=17), and Hom-NMyc (n=8). No significant differences were observed (ANOVA).

**Figure S3.**
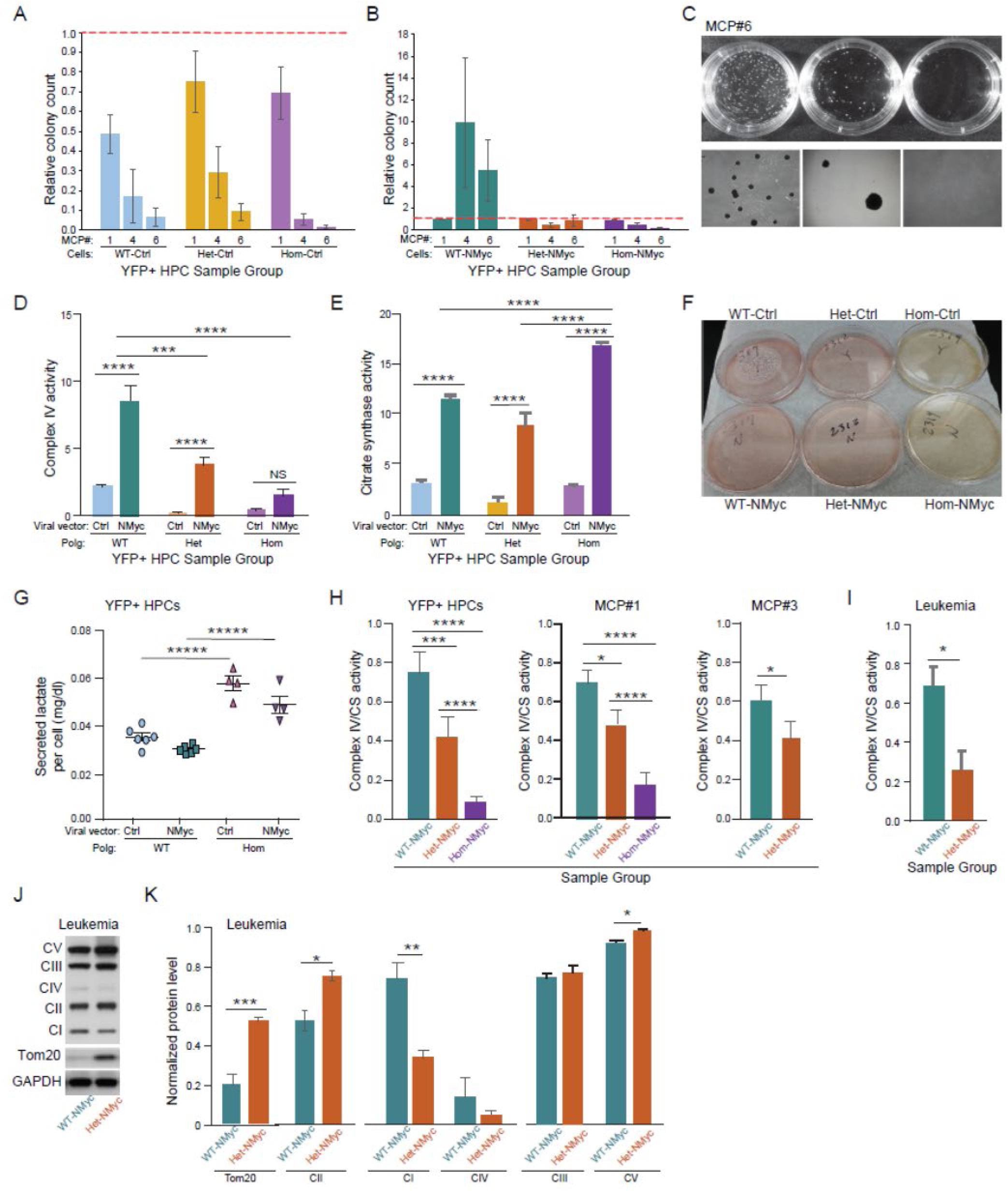
Supplement to Figure 3. **(A-B)** Colony numbers were assessed in serial methylcellulose culture passages (MCP#) at MCP#1, MCP#4, and MCP#6 across different groups and then normalized to the number of WT-NMyc cells at MCP#1. **(A)** Bar graphs showing results for the control groups (WT-Ctrl, Het-Ctrl, Hom-Ctrl) show loss of colonies by MCP#6. **(B)** Bar graphs showing results for the NMyc expression HPC groups (WT-NMyc, Het-NMyc, Hom-NMyc) show persistence of WT-NMyc colonies at MCP#6. Sample numbers for **(A)** and **(B)**: n=13-14 biologic replicates per group at MCP#1; n=9-10 biologic replicates per group at MCP#4; n=8-9 biologic replicates per group at MCP#6. **(C)** Photographs of plates with colonies derived from NMyc expressing HPCS at MCP#6 (upper panel) and corresponding light microscopy images (lower panel). **(D, E)** Complex IV **(D)** and citrate synthase **(E)** activities were measured using sorted HPCs (pooled from two mice for each genotype). Results from a representative experiment (n=4 technical replicates for each group) are shown. ***P<0.005, ****P<0.00001 (ANOVA). (**F**) At MCP#1 (after 1 week in culture), the methylcellulose media in cultures containing Hom-Ctrl and Hom-Nmyc HPCs changed color, signifying acidification of the media. **(G)** Secreted lactate was measured in WT and Hom HPCs, with or without NMyc expression. WT-Ctrl (n=6), WT-NMyc (n=6), Het-Ctrl (n=4), Hom-NMyc (n=4) *****P<0.0005. (Tukey’s multiple comparisons test). **(H)** Bar graph shows the ratio of complex IV to citrate synthase activity (“Normalized complex IV activity”) using sorted YFP+ HPCs (pooled from two mice for each genotype) before (i.e., YFP+ HPC) and after culturing in methylcellulose (i.e., MCP#1, MCP#3). Results from a representative experiment (n=4 technical replicates for each group) are shown. **(I)** Normalized complex IV activity was measured using lysates prepared from leukemia-infiltrated spleens from WT-NMyc (n=3) and Het-NMyc (3) mice. All significant differences were measured with an ANOVA, *P<0.05, ***P<0.005, ****P<0.001. **(J-L)** Lysates were prepared from leukemia-infiltrated spleens of WT-NMyc (n=3) and Het-NMyc (n=3) mice. Representative immunoblots **(J)** using antibodies against labile OXPHOS components (complexes I–V), Tom20 and GAPDH. Graphs showing normalized protein levels (mean + s.e.m.) are shown **(K)** *P<0.05, **P<0.01, ***P<0.005 (Student’s *t*-test).

**Figure S4.**
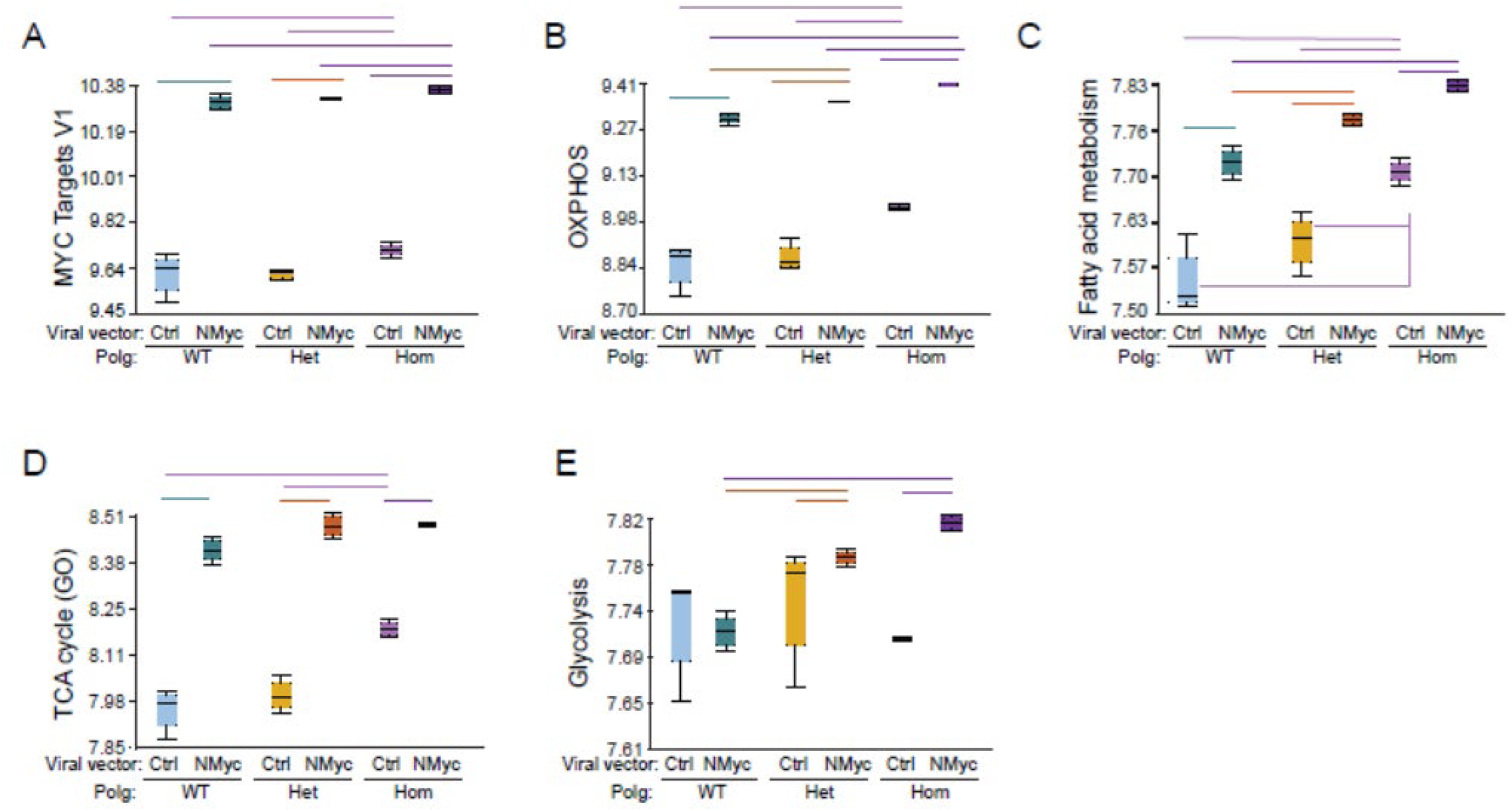
Supplement to Figure 4. **(A-E)** Mean pathway expression values across 6 HPC groups for the following genesets: **(A)** MYC Targets Version 1 (v1, Hallmark) **(B)** OXPHOS (Hallmark), **(C)** fatty acid metabolism (Hallmark), **(D)** Genes related to TCA cycle (GO)**, and (E)** Glycolysis (Hallmark). WT-Ctrl (n=5), WT-NMyc (n=5), Het-Ctrl (n=5), Het-NMyc (n=5), Hom-Ctrl (n=5), and Hom-NMyc (n=5). Results of ANOVA are provided in **Table S2**.

**Figure S5.**
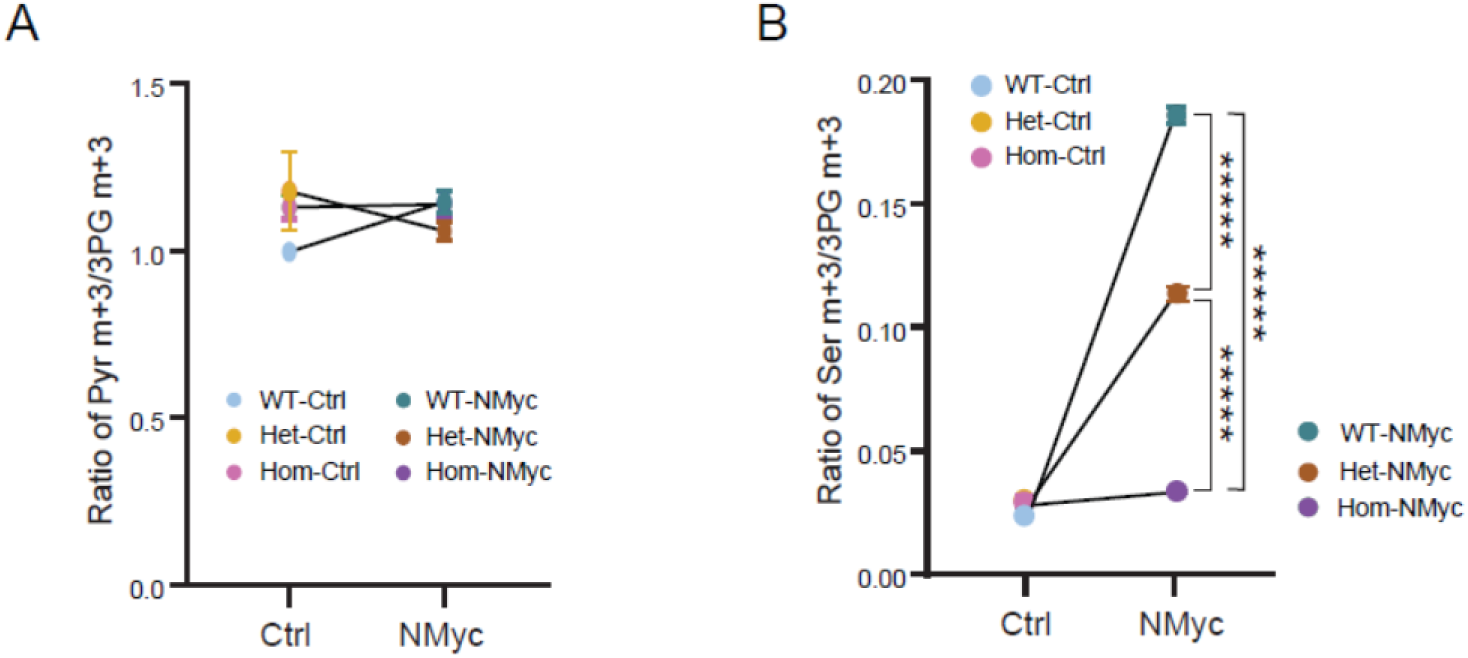
Supplement to Figure 5. **(A)** Ratio of Pyr m+3/3PG m+3 was measured in both the control and NMyc groups. **(B)** Ratio of Ser m+3/3PG m+3 was measured for all genotypes, *****P<0.0005 (Student’s *t*-test). WT-Ctrl (n=5), WT-NMyc (n=5), Het-Ctrl (n=6), Het-NMyc (n=4), Hom-Ctrl (n=5), Hom-NMyc (n=6).

**Figure S6.**
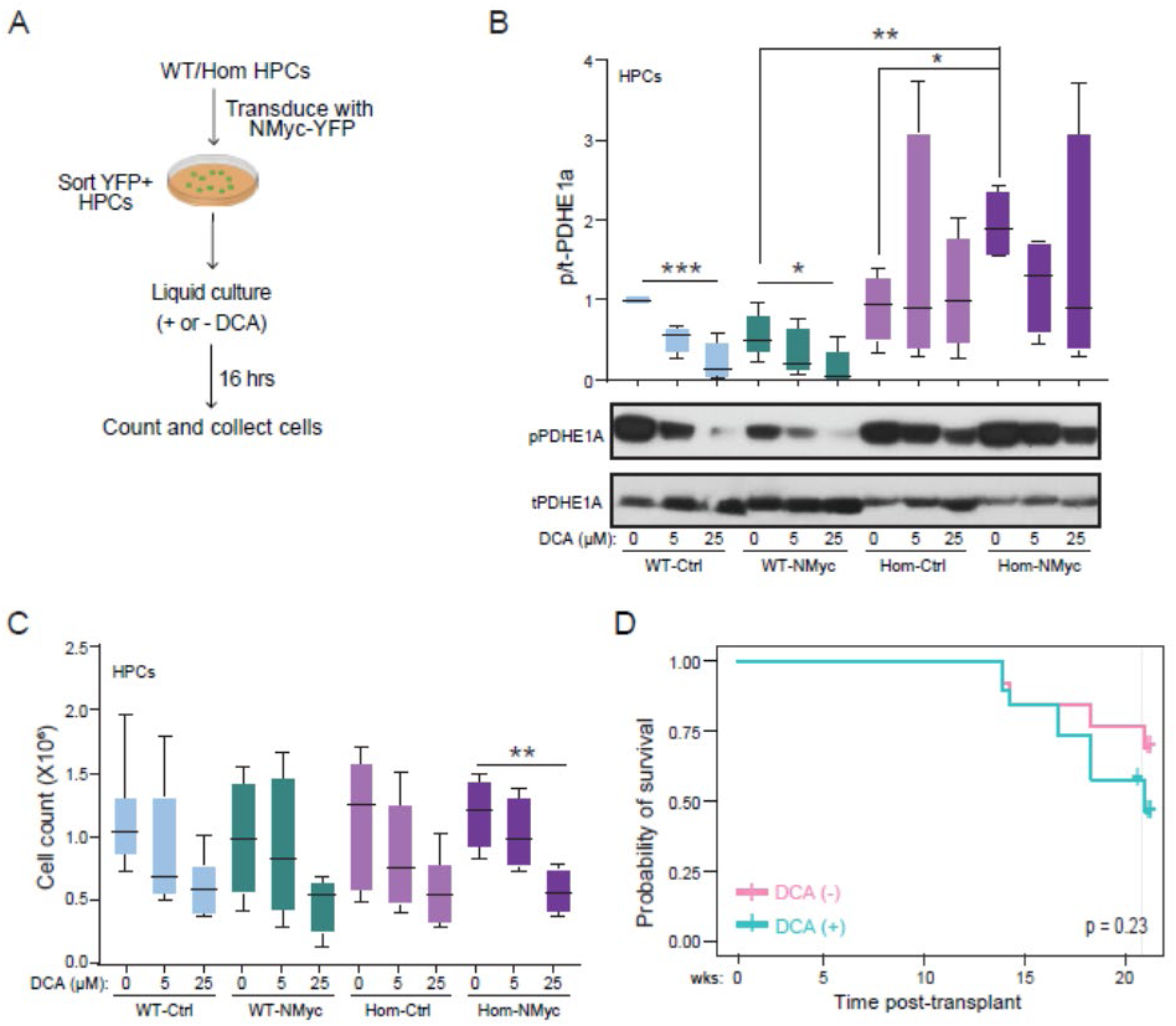
Supplement to Figure 6. **(A)** Schematic of the in vitro experiments to determine the effects of dichloroacetate (DCA) treatment on the growth of HPCs cultured in liquid media for 16 h. (**B**) Immunoblot analysis was performed using lysates prepared from control and NMyc-expressing WT, Het, and Hom samples after DCA treatment at 0, 5, or 25 µM (bottom). The normalized protein level of phosphorylated PDHE1A (p-PDHE1A) by t-PDHE1A was presented (top). (**C**) Cells were counted after 16-h DCA treatment using 0-µM, 5-µM or 25-µM DCA. n=4-6 biologic replicates per condition per group Statistical significance for **(B, C)** was determined by ANOVA, *P<0.05, **P<0.01, ***P<0.005. **(D)** Kaplan-Meier survival curve for Hom-NMyc mice treated with or without DCA.

## REFERENCES

1. I. Martinez-Reyes, N. S. Chandel, Cancer metabolism: looking forward. Nat Rev Cancer 21, 669–680 (2021).

2. S. Papadaki, A. Magklara, Regulation of Metabolic Plasticity in Cancer Stem Cells and Implications in Cancer Therapy. Cancers (Basel) 14, (2022).

3. P. E. Porporato, N. Filigheddu, J. M. B. Pedro, G. Kroemer, L. Galluzzi, Mitochondrial metabolism and cancer. Cell Res 28, 265–280 (2018).

4. K. Vasan, M. Werner, N. S. Chandel, Mitochondrial Metabolism as a Target for Cancer Therapy. Cell Metab 32, 341–352 (2020).

5. P. K. Kopinski, L. N. Singh, S. Zhang, M. T. Lott, D. C. Wallace, Mitochondrial DNA variation and cancer. Nat Rev Cancer 21, 431–445 (2021).

6. A. L. M. Smith, J. C. Whitehall, L. C. Greaves, Mitochondrial DNA mutations in ageing and cancer. Mol Oncol 16, 3276–3294 (2022).

7. J. A. Petros et al., mtDNA mutations increase tumorigenicity in prostate cancer. Proc Natl Acad Sci U S A 102, 719–724 (2005).

8. M. F. Marusich et al., Expression of mtDNA and nDNA encoded respiratory chain proteins in chemically and genetically-derived Rho0 human fibroblasts: a comparison of subunit proteins in normal fibroblasts treated with ethidium bromide and fibroblasts from a patient with mtDNA depletion syndrome. Biochim Biophys Acta 1362, 145–159 (1997).

9. D. C. Wallace, Mitochondrial diseases in man and mouse. Science 283, 1482–1488 (1999).

10. A. S. Tan et al., Mitochondrial genome acquisition restores respiratory function and tumorigenic potential of cancer cells without mitochondrial DNA. Cell Metab 21, 81–94 (2015).

11. L. F. Dong et al., Horizontal transfer of whole mitochondria restores tumorigenic potential in mitochondrial DNA-deficient cancer cells. Elife 6, (2017).

12. P. A. Gammage, C. Frezza, Mitochondrial DNA: the overlooked oncogenome? BMC Biol 17, 53 (2019).

13. C. Welter, G. Kovacs, G. Seitz, N. Blin, Alteration of mitochondrial DNA in human oncocytomas. Genes Chromosomes Cancer 1, 79–82 (1989).

14. K. Polyak et al., Somatic mutations of the mitochondrial genome in human colorectal tumours. Nat Genet 20, 291–293 (1998).

15. E. R. Mardis et al., Recurring mutations found by sequencing an acute myeloid leukemia genome. N Engl J Med 361, 1058–1066 (2009).

16. F. Damm et al., Prognostic implications and molecular associations of NADH dehydrogenase subunit 4 (ND4) mutations in acute myeloid leukemia. Leukemia 26, 289–295 (2012).

17. Y. S. Ju et al., Origins and functional consequences of somatic mitochondrial DNA mutations in human cancer. Elife 3, (2014).

18. Y. Yuan et al., Comprehensive molecular characterization of mitochondrial genomes in human cancers. Nat Genet 52, 342–352 (2020).

19. M. Picard et al., Progressive increase in mtDNA 3243A>G heteroplasmy causes abrupt transcriptional reprogramming. Proc Natl Acad Sci U S A 111, E4033–4042 (2014).

20. J. S. Park et al., A heteroplasmic, not homoplasmic, mitochondrial DNA mutation promotes tumorigenesis via alteration in reactive oxygen species generation and apoptosis. Hum Mol Genet 18, 1578–1589 (2009).

21. G. C. Kujoth et al., Mitochondrial DNA mutations, oxidative stress, and apoptosis in mammalian aging. Science 309, 481–484 (2005).

22. A. Trifunovic et al., Premature ageing in mice expressing defective mitochondrial DNA polymerase. Nature 429, 417–423 (2004).

23. J. E. Kolesar et al., Defects in mitochondrial DNA replication and oxidative damage in muscle of mtDNA mutator mice. Free Radic Biol Med 75, 241–251 (2014).

24. T. Mito et al., Mitochondrial DNA mutations in mutator mice confer respiration defects and B-cell lymphoma development. PLoS One 8, e55789 (2013).

25. H. Kawagoe, A. Kandilci, T. A. Kranenburg, G. C. Grosveld, Overexpression of N-Myc rapidly causes acute myeloid leukemia in mice. Cancer Res 67, 10677–10685 (2007).

26. A. Astolfi et al., MYCN is a novel oncogenic target in pediatric T-cell acute lymphoblastic leukemia. Oncotarget 5, 120–130 (2014).

27. F. Li et al., Myc stimulates nuclearly encoded mitochondrial genes and mitochondrial biogenesis. Mol Cell Biol 25, 6225–6234 (2005).

28. F. R. Dejure, M. Eilers, MYC and tumor metabolism: chicken and egg. EMBO J 36, 3409–3420 (2017).

29. H. Hirvonen, V. Hukkanen, T. T. Salmi, T. T. Pelliniemi, R. Alitalo, L-myc and N-myc in hematopoietic malignancies. Leuk Lymphoma 11, 197–205 (1993).

30. W. Zhang, H. Cui, L. J. Wong, Comprehensive one-step molecular analyses of mitochondrial genome by massively parallel sequencing. Clin Chem 58, 1322–1331 (2012).

31. K. D. Maclaine, K. A. Stebbings, D. A. Llano, J. C. Havird, The mtDNA mutation spectrum in the PolG mutator mouse reveals germline and somatic selection. BMC Genom Data 22, 52 (2021).

32. A. Ameur et al., Ultra-Deep Sequencing of Mouse Mitochondrial DNA: Mutational Patterns and Their Origins. Plos Genetics 7, (2011).

33. M. L. Chen et al., Erythroid dysplasia, megaloblastic anemia, and impaired lymphopoiesis arising from mitochondrial dysfunction. Blood 114, 4045–4053 (2009).

34. G. L. Norddahl et al., Accumulating mitochondrial DNA mutations drive premature hematopoietic aging phenotypes distinct from physiological stem cell aging. Cell Stem Cell 8, 499–510 (2011).

35. X. Li-Harms et al., Mito-protective autophagy is impaired in erythroid cells of aged mtDNA-mutator mice. Blood 125, 162–174 (2015).

36. G. Hoxhaj, B. D. Manning, The PI3K-AKT network at the interface of oncogenic signalling and cancer metabolism. Nat Rev Cancer 20, 74–88 (2020).

37. L. Zuurbier et al., The significance of PTEN and AKT aberrations in pediatric T-cell acute lymphoblastic leukemia. Haematologica 97, 1405–1413 (2012).

38. A. Stengel et al., The impact of TP53 mutations and TP53 deletions on survival varies between AML, ALL, MDS and CLL: an analysis of 3307 cases. Leukemia 31, 705–711 (2017).

39. I. S. Jerchel et al., RAS pathway mutations as a predictive biomarker for treatment adaptation in pediatric B-cell precursor acute lymphoblastic leukemia. Leukemia 32, 931–940 (2018).

40. Y. Fukuda, et al., Upregulated heme biosynthesis, an exploitable vulnerability in MYCN-driven leukemogenesis. JCI Insight 2, (2017).

41. M. Guha, N. G. Avadhani, Mitochondrial retrograde signaling at the crossroads of tumor bioenergetics, genetics and epigenetics. Mitochondrion 13, 577–591 (2013).

42. R. A. Hardie et al., Mitochondrial mutations and metabolic adaptation in pancreatic cancer. Cancer Metab 5, 2 (2017).

43. J. Zhang et al., Measuring energy metabolism in cultured cells, including human pluripotent stem cells and differentiated cells. Nat Protoc 7, 1068–1085 (2012).

44. A. Subramanian et al., Gene set enrichment analysis: a knowledge-based approach for interpreting genome-wide expression profiles. Proc Natl Acad Sci U S A 102, 15545–15550 (2005).

45. A. Liberzon et al., Molecular signatures database (MSigDB) 3.0. Bioinformatics 27, 1739–1740 (2011).

46. A. Liberzon et al., The Molecular Signatures Database (MSigDB) hallmark gene set collection. Cell Syst 1, 417–425 (2015).

47. T. McFate et al., Pyruvate dehydrogenase complex activity controls metabolic and malignant phenotype in cancer cells. J Biol Chem 283, 22700–22708 (2008).

48. G. Sutendra et al., A nuclear pyruvate dehydrogenase complex is important for the generation of acetyl-CoA and histone acetylation. Cell 158, 84–97 (2014).

49. B. L. Woolbright, G. Rajendran, R. A. Harris, J. A. Taylor, 3rd, Metabolic Flexibility in Cancer: Targeting the Pyruvate Dehydrogenase Kinase:Pyruvate Dehydrogenase Axis. Mol Cancer Ther 18, 1673–1681 (2019).

50. X. Wang, X. Shen, Y. Yan, H. Li, Pyruvate dehydrogenase kinases (PDKs): an overview toward clinical applications. Biosci Rep 41, (2021).

51. L. Yang et al., NAD(+) dependent UPR(mt) activation underlies intestinal aging caused by mitochondrial DNA mutations. Nat Commun 15, 546 (2024).

52. K. X. Liang et al., Activation of Neurotoxic Astrocytes Due to Mitochondrial Dysfunction Triggered by POLG Mutation. Int J Biol Sci 20, 2860–2880 (2024).

53. K. A. Olson, J. C. Schell, J. Rutter, Pyruvate and Metabolic Flexibility: Illuminating a Path Toward Selective Cancer Therapies. Trends Biochem Sci 41, 219–230 (2016).

54. E. D. Michelakis, L. Webster, J. R. Mackey, Dichloroacetate (DCA) as a potential metabolic-targeting therapy for cancer. Br J Cancer 99, 989–994 (2008).

55. W. Y. Sanchez et al., Dichloroacetate inhibits aerobic glycolysis in multiple myeloma cells and increases sensitivity to bortezomib. Br J Cancer 108, 1624–1633 (2013).

56. L. Yu et al., Modeling the Genetic Regulation of Cancer Metabolism: Interplay between Glycolysis and Oxidative Phosphorylation. Cancer Res 77, 1564–1574 (2017).

57. G. Cannino, F. Ciscato, I. Masgras, C. Sanchez-Martin, A. Rasola, Metabolic Plasticity of Tumor Cell Mitochondria. Front Oncol 8, 333 (2018).

58. D. Jia et al., Elucidating cancer metabolic plasticity by coupling gene regulation with metabolic pathways. Proc Natl Acad Sci U S A 116, 3909–3918 (2019).

59. N. Niu, J. Ye, Z. Hu, J. Zhang, Y. Wang, Regulative Roles of Metabolic Plasticity Caused by Mitochondrial Oxidative Phosphorylation and Glycolysis on the Initiation and Progression of Tumorigenesis. Int J Mol Sci 24, (2023).

60. S. Delaunay et al., Mitochondrial RNA modifications shape metabolic plasticity in metastasis. Nature 607, 593–603 (2022).

61. D. C. Wallace, Mitochondria and cancer. Nat Rev Cancer 12, 685–698 (2012).

62. P. Y. Wang et al., Inhibiting mitochondrial respiration prevents cancer in a mouse model of Li-Fraumeni syndrome. J Clin Invest 127, 132–136 (2017).

63. H. A. Prag, M. P. Murphy, mtDNA mutations help support cancer cells. Nat Cancer 1, 941–942 (2020).

64. R. Rathore, C. R. Schutt, B. A. Van Tine, PHGDH as a mechanism for resistance in metabolically-driven cancers. Cancer Drug Resist 3, 762–774 (2020).

65. M. J. Holness, M. C. Sugden, Regulation of pyruvate dehydrogenase complex activity by reversible phosphorylation. Biochem Soc Trans 31, 1143–1151 (2003).

66. R. A. Cairns, I. S. Harris, T. W. Mak, Regulation of cancer cell metabolism. Nat Rev Cancer 11, 85–95 (2011).

67. M. G. Vander Heiden, L. C. Cantley, C. B. Thompson, Understanding the Warburg effect: the metabolic requirements of cell proliferation. Science 324, 1029–1033 (2009).

68. P. S. Ward, C. B. Thompson, Metabolic reprogramming: a cancer hallmark even warburg did not anticipate. Cancer Cell 21, 297–308 (2012).

69. R. A. Irizarry et al., Exploration, normalization, and summaries of high density oligonucleotide array probe level data. Biostatistics 4, 249–264 (2003).

70. P. A. Srere, in Methods in Enzymology. (Academic Press, 1969), vol. 13, pp. 3–11.

71. I. F. Robey, N. K. Martin, Bicarbonate and dichloroacetate: evaluating pH altering therapies in a mouse model for metastatic breast cancer. BMC Cancer 11, 235 (2011).

72. D. P. Harrington, T. R. Fleming, A class of rank test procedures for censored survival data. Biometrika 69, 553–566 (1982).

73. R. J. Gray, A Class of K-Sample Tests for Comparing the Cumulative Incidence of a Competing Risk. The Annals of Statistics 16, 1141–1154 (1988).

74. E. Demidenko, Mixed Models: Theory and Applications with R. John Wiley & Sons. Aug 26, 2013

